# Epoxidized graphene grid for high-throughput high-resolution cryoEM structural analysis

**DOI:** 10.1101/2021.11.17.468963

**Authors:** Junso Fujita, Fumiaki Makino, Haruyasu Asahara, Maiko Moriguchi, Shota Kumano, Itsuki Anzai, Jun-ichi Kishikawa, Yoshiharu Matsuura, Takayuki Kato, Keiichi Namba, Tsuyoshi Inoue

## Abstract

Many specimens suffer from low particle density and/or preferred orientation in cryoEM specimen grid preparation, making data collection and structure determination time consuming. We developed an epoxidized graphene grid (EG-grid) that effectively immobilizes protein particles by applying an oxidation reaction using photoactivated ClO_2_^•^ and further chemical modification. The particle density and orientation distribution are both dramatically improved, having enabled us to reconstruct the density map of GroEL and glyceraldehyde 3-phosphate dehydrogenase (GAPDH), at 1.99 and 2.16 Å resolution from only 504 and 241 micrographs, respectively. A low concentration sample solution of 0.1 mg ml^−1^ was sufficient to reconstruct a 3.10 Å resolution density map of SARS-CoV-2 spike protein from 1,163 micrographs. The density maps of V_1_-ATPase, β-galactosidase, and apoferritin were also reconstructed at 3.03, 1.81, and 1.29 Å resolution, respectively. These results indicate that the EG-grid will be a powerful tool for high-throughput cryoEM data collection to accelerate high-resolution structural analysis of biological macromolecules.

## Introduction

Despite the recent improvements in hardware and software for electron cryomicroscopy (cryoEM) single particle image analysis for high-throughput and high-resolution structure determination of biological macromolecules, specimen grid preparation still remains a bottleneck due to various factors. In single particle analysis (SPA), protein particles must be well dispersed in aqueous solution to be flash-frozen and embedded in thin vitreous ice film formed within micron-sized holes on a few 10s nm thick carbon film laid on a metal grid substrate^1^. Automated vitrification devices have been developed^2–5^, and various efforts have been made to produce good grids with well-dispersed particles and without much sample loss by denaturation^4–8^. In general, proteins often localize to the air-water interface and denature, resulting in poor particle density and preferred orientation and thereby making high-resolution 3D reconstruction inefficient^9^. Due to these problems, optimizing the conditions for cryoEM grid preparation requires much time and effort^10,11^. To overcome these problems, thin support films, such as graphene, have been used to change the physical properties of grid surface^12–15^, but in cases where preferred orientation still persists, the grid needs be tilted for data collection^16,17^. Although graphene itself is unsuitable support due to its hydrophobicity, it can be hydrophilized by conventional glow discharge or chemical modification^18^. Graphene oxide synthesized from graphite using modified Hummers’ methods^19,20^ is one of the most common derivatives and easy to obtain at low cost, but its fragmented nature tends to produce non-uniform coverage of the grid surface.

Graphene synthesized by chemical vapor deposition on metal substrates, such as Cu, forms a large uniform sheet suitable to cover the entire surface of metal grids, and there have been some cryoEM studies using functionalized graphene grids^13,14^. A graphene oxide grid modified with Nα, Nα-dicarboxymethyllysine (NTA) has been designed to capture His-tagged proteins^21,22^. Introduction of a SpyTag consisting of 13 amino acid residues into a target protein or the introduction of Cys residues has been reported also successful for cryoEM analysis by making proteins tightly bound to a graphene oxide grid^23^. These affinity grids helped to minimize the exposure of proteins to the air-water interface and improved the data quality, but introduction of specific tags to the target protein is required. One potential solution recently proposed was the introduction of amino groups and polyethylene glycol linkers on graphene oxide, which kept most of proteins away from both the air-water interface and graphene oxide surface and improved particle distribution and orientation^24^.

It would be more efficient and useful if we can immobilize any target proteins directly on the graphene surface and keep them intact. We now developed an epoxidized graphene grid (EG-grid) that effectively immobilizes protein particles by applying an oxidation reaction using photoactivated ClO_2_^•^ and further chemical modification. We have recently discovered a useful chemical reaction using photoactivated ClO_2_^•^ gas as a strong oxidant; it can convert a gaseous fuels of methane into liquid fuels of methanol and formic acid without producing CO_2_^25^ and can also make surface oxidation modification of polymers, such as polypropylene^26^. We applied this oxidation reaction to introduce hydroxy groups into a basal plane of graphene surface and further introduced epoxy groups by epichlorohydrin (ECH) treatment, which has been used for functionalization of graphene oxide^27,28^. This EG-grid effectively immobilized proteins without tags so that they stayed on the surface even after washing with buffer. We tested it by collecting cryoEM data of GroEL, glyceraldehyde 3-phosphate dehydrogenase (GAPDH), β-galactosidase, and apoferritin, and many of them showed high particle density even with relatively low protein concentrations of the sample solutions, enabling effective data collection just by one grid preparation each. The resolutions of the final 3D maps of GroEL, GAPDH, β-galactosidase, and apoferritin reached 1.99, 2.16, 1.81, and 1.29 Å, respectively. Especially with GroEL and GAPDH, only 504 and 241 micrographs were needed to achieve such high resolutions. We also demonstrated effective data collection of SARS-CoV-2 spike protein on the EG-grid with a sample solution at a protein concentration of only 0.1 mg ml^−1^, and the resolution of the reconstructed map reached 3.10 Å. V_1_-ATPase was also tested as a sample of preferred orientation, and the map resolution was improved from ~5 Å to 3.03 Å by much more uniform angular distribution. We believe this EG-grid will become a powerful tool for accelerating high-resolution cryoEM structure determination.

## Results

### Characterization of ClO_2_^•^ oxidized graphene

The EG-grid is produced in three steps as shown in a schematic illustration of the fabrication process (Fig. 1a). First, a CVD graphene layer is put on a Quantifoil Au grid as previously reported^13^ with some modification. Then, the surface of graphene is oxidized by photoactivated ClO_2_^•^ treatment for 10 min to introduce hydroxy groups (see Methods for detail and Supplementary Video 1). The following reaction with ECH in the third step introduces epoxy groups on the surface of graphene. The epoxy groups are expected to form covalent bonds with nucleophilic groups such as amino, hydroxy, or thiol groups distributed on the surface of proteins. The hydrophilicity of graphene grids was evaluated by the water contact angle (WCA) measurement. The WCA of graphene on the Au grid decreased from 74° to 60° by the ClO_2_^•^ oxidation treatment (Fig. 1a, left panels). Furthermore, the introduction of epoxy groups slightly decreased the hydrophilicity (65°).

**Fig. 1.**
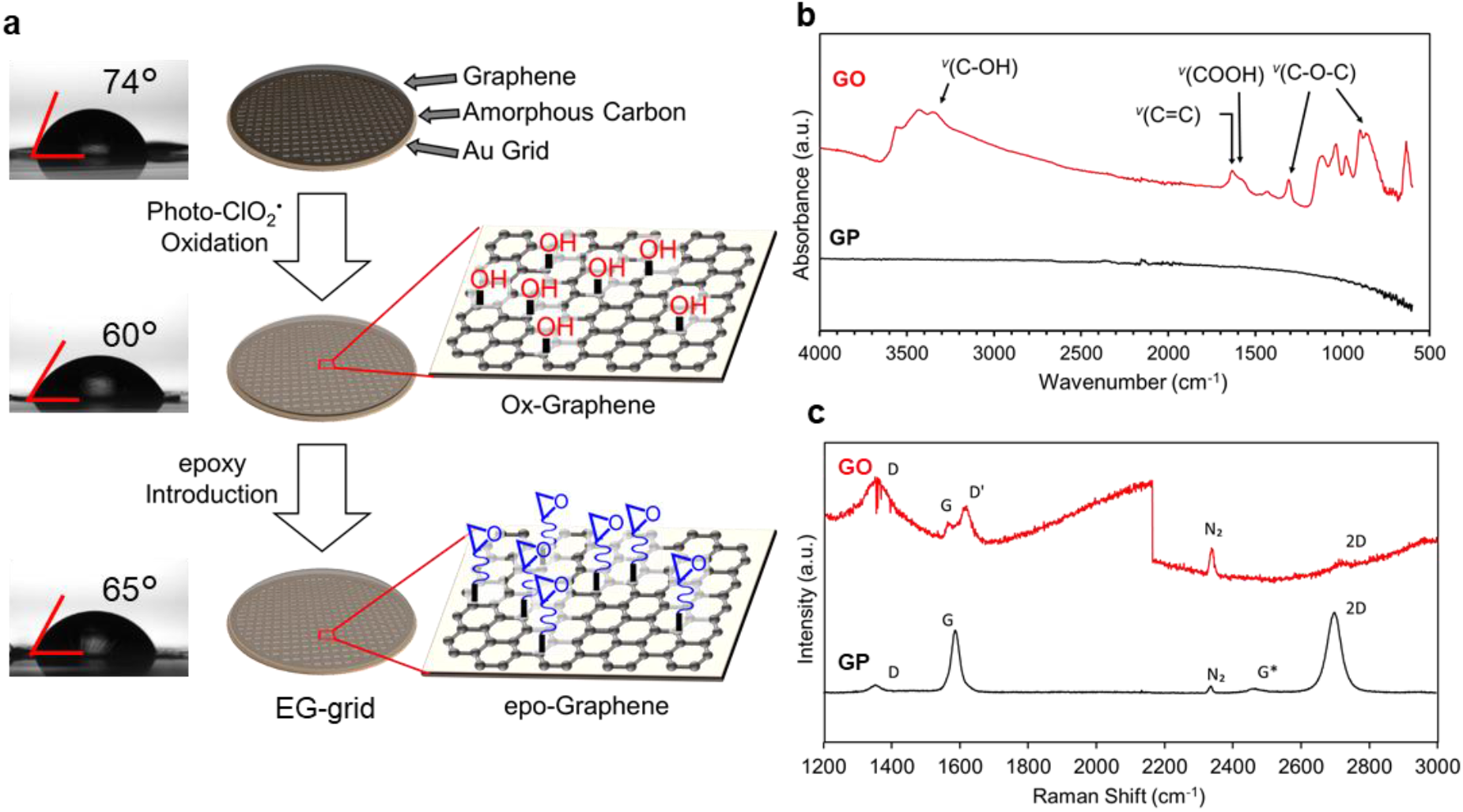
Fabrication process of the EG-grid and characterization of the graphene surface. **a**, Water contact angle measurement and schematic illustration of EG-grid preparation by oxidation and functionalization of graphene on the Quantifoil Au grid. **b**, IR spectra of pristine graphene (GP: black line) and oxidized graphene after 10 min ClO_2_^•^ treatment (GO: red line). **c**, Raman spectra of GP (black line) and GO (red line) on silicon wafer.

To confirm the oxidation state of the graphene surface, we performed infrared (IR) and Raman spectral measurements and analyses. While pristine graphene (GP) was IR inactive as shown in Fig. 1b, graphene oxidized by ClO_2_^•^ (GO) showed several characteristic peaks such as those at 3400 cm^-1^, ^*v*^(C-OH); 1620 cm^-1^, ^*v*^(C=C); and 1300 cm^-1^, 850 cm^-1^, ^*v*^(C-O-C), suggesting that oxygen-containing functional groups were successfully introduced to the graphene surface. The Raman spectra of GP and GO are shown in Fig. 1c, in which GP showed two main peaks: G (1580 cm^-1^) and 2D (2690 cm^-1^), and their intensity ratio (I_2D_/I_G_) was about 1.8, proving the high quality of single layer graphene. After oxidation, the D and D’ peaks appeared whereas the 2D peak nearly diminished. These features of the Raman spectra indicated the formation of sp^3^-carbon by the introduction of functional groups to graphene basal plane.

To investigate the effect of oxidation on the chemical state of the graphene surface more comprehensively, the chemical compositions of the functional groups were determined by X-ray photoelectron spectroscopy (XPS) measurement. Graphene was transferred onto a silicon wafer and then oxidized either by plasma or photoactivated ClO_2_^•^ treatment. After plasma treatment, the composition ratios of the C-O groups (corresponding to the peak at 286.0 eV) and the C=O groups (corresponding to the peak at 287.8 eV) were 22.5% and 29.2%, respectively (Supplementary Fig. 1). On the other hand, in the ClO_2_^•^-treated graphene, they were 28.4% (C-O) and 8.5% (C=O), respectively. Generally, the increase of C=O ratio means the cleavage of the C=C bond of graphene. The increase of C-O ratio indicated that the hydroxy groups were introduced dominantly to the basal plane in the ClO_2_^•^-treated graphene rather than the cleavage of the C=C bond in the plasma-treated graphene, in agreement with the IR measurement.

To confirm the introduction of epoxy groups and formation of covalent bonds between the epoxy and amino groups, we used 1H,1H-undecafluorohexylamine (UFHA) because fluorine peaks can be clearly observed in XPS measurement (Supplementary Fig. 2). Only carbon and oxygen were detected with non-oxidized graphene (black line). No fluorine was detected with UFHA-treated oxidized graphene and ECH-/UFHA-treated graphene (red and green lines), while clear fluorine peak appeared with UFHA-treated EG-grid (purple line). These results indicated that UFHA was covalently bonded to epoxy group on the graphene surface.

### Protein binding activity of the EG-grid surface

To investigate the protein binding activity of the EG-grid, we immobilized GroEL from *Escherichia coli* as a benchmark protein because the resolution of the cryoEM 3D reconstruction of this protein has been limited to 3.26 Å possibly by limited number of particles due to elimination of damaged particle images and by preferred orientation towards its end-on views^29^. A 3 μl solution of GroEL at a protein concentration of 0.3 mg ml^−1^ was applied on an EG-grid, and the protein solution was blotted after 5 min incubation. The grid was washed three times with 5 μl of buffer each, and then the grid was negatively stained with uranyl acetate. Many GroEL particles remained on the EG-grid and were packed densely (Supplementary Fig. 3). We also prepared a grid in the same way except skipping the epoxidation process by ECH and observed no or very small number of GroEL particles, which indicates that the EG-grid strongly adsorbed the protein particles.

### CryoEM image analysis of GroEL on EG-grid

For cryoEM image analysis of GroEL, 3 μl of 1.5 mg ml^−1^ GroEL solution was loaded onto the EG-grid and allowed to stand for 5 min at room temperature, followed by blotting and plunge freezing using Vitrobot (Thermo Fisher Scientific). A short blotting time around 1.0–1.5 s usually works well for getting appropriate ice thickness on the EG-grid. The grid was then observed using an electron cryomicroscope (cryoTEM) equipped with a cold field emission gun and an Ω-type in-column energy filter (JEM-Z300FSC (CRYO ARM 300), JEOL) operated at 300 kV, and cryoEM images were recorded on a K3 direct electron detector (Gatan, Inc.). The micrographs showed more densely packed GroEL particles with much more side view orientations (Fig. 2a) than cases using standard Quantifoil grids^30^. We collected a dataset at a nominal magnification of 60,000× (referred to as GroEL-EG). For comparison with a more conventional grid preparation method, we applied the same GroEL solution to a Quantifoil grid covered with graphene hydrophilized by glow discharge. We collected a dataset with the same cryoTEM and settings (referred to as GroEL-Gl). The results showed that the density of the GroEL particles was much lower than that on the EG-grid, with more top view orientations (Fig. 2b). To confirm whether the difference in the orientation distribution between these grids was caused by the EG-grid, we collected another dataset from a different glow discharged graphene grid (referred to as GroEL-Gh). Although the grid was washed with the buffer after loading 1.0 mg ml^−1^ protein solution, densely packed and side-view oriented particles were observed similarly to GroEL-EG in some area of the grid (Fig. 2c).

**Fig. 2.**
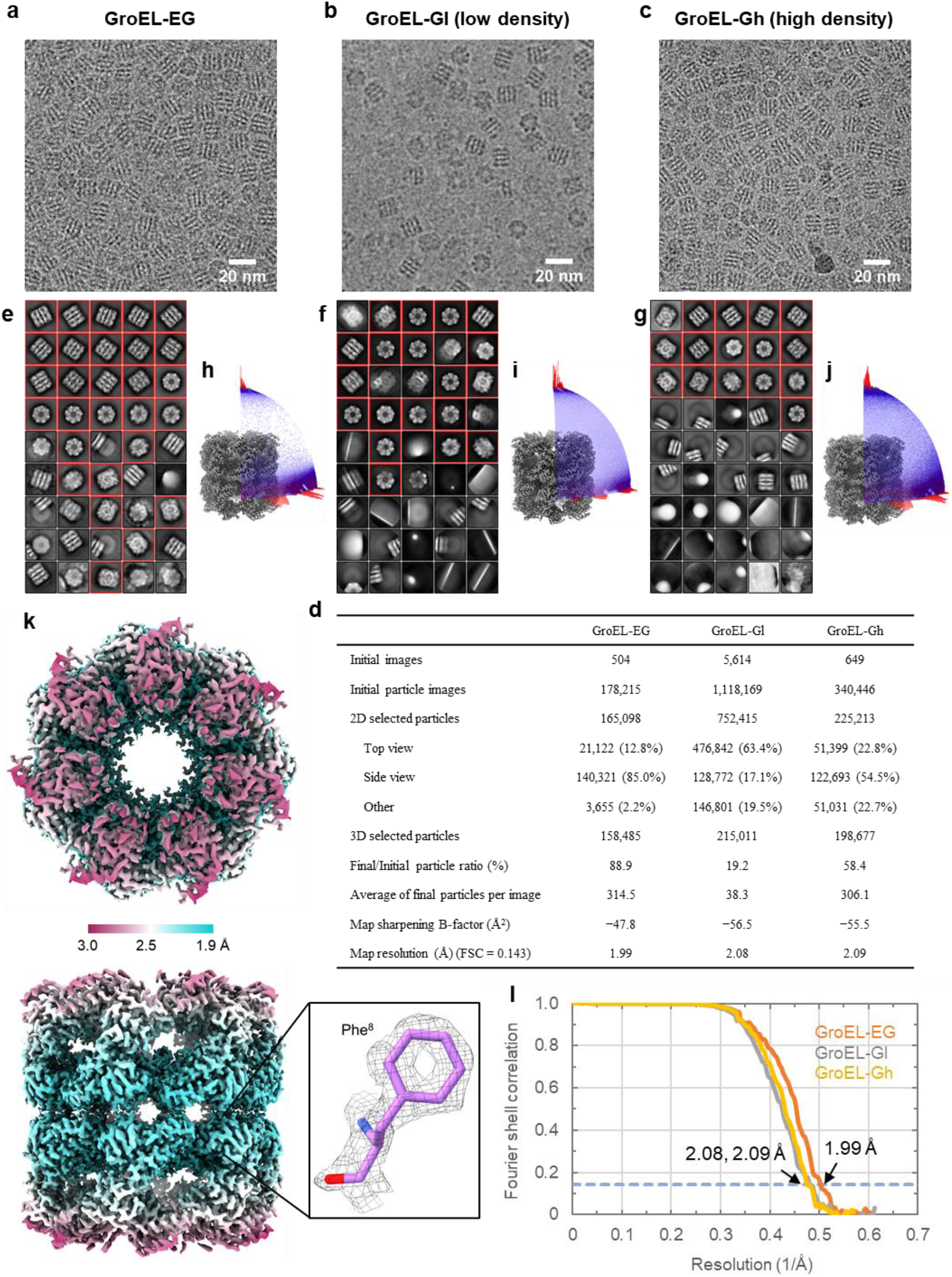
CryoEM image analyses of GroEL on the EG-grid and glow discharged graphene grid. **a**–**c**, Typical cryoEM images of GroEL on the EG-grid (GroEL-EG) (**a**), and GroEL packed with low density (GroEL-Gl) (**b**) and high density (GroEL-Gh) (**c**) on the different glow discharged graphene grids. **d**, Selected statistics of cryoEM image processing of the three datasets. **e**–**g**, Selected 2D class averages from the GroEL-EG (**e**), GroEL-Gl (**f**), and GroEL-Gh (**g**) dataset. The class averages are aligned in descending order of particle numbers from left to right and top to bottom. The classes selected for the following analysis are indicated with red boxes. **h**–**j**, Angular distribution of particles used in the final refinement for GroEL-EG (**h**), GroEL-Gl (**i**), and GroEL-Gh (**j**) dataset. Final 3D maps are also shown for reference. **k**, Two orthogonal views of the final 3D map from the GroEL-EG dataset: top view, upper panel; and side view, lower panel. The local resolution distribution is colored as in the color bar. The inset shows the density map and the fitted model (PDB: 1SX3) around Phe 8. **l**, The FSC curves for the final maps of the GroEL-EG, GroEL-Gl, and GroEL-Gh dataset. The dashed blue line indicates the FSC = 0.143 criterion.

The three datasets were processed with Relion 3.1. Images of 158,485, 215,011, and 198,677 particles were used for final reconstructions from 504, 5,614, and 649 micrographs, resulting in 3D maps with resolutions of 1.99, 2.08, and 2.09 Å for the GroEL-EG, GroEL-Gl, and GroEL-Gh dataset, respectively (Fig. 2d and Supplementary Table 1). The low particle density and top-preferred orientation in the GroEL-Gl dataset resulted in a slightly lower resolution than that of the GroEL-EG dataset even from more than 10 times movie images of the latter (Fig. 2d-f,h,i). We counted the average number of particles per image contributing to the final reconstruction (NPPI-final), which would reflect the efficiency of data collection. For the GroEL-EG and GroEL-Gh dataset, 315 and 306 particles/image were used for final reconstruction, respectively, while only 38 particles/image were used for the GroEL-Gl dataset (Fig. 2d and Supplementary Table 1).

Although both the GroEL-EG and GroEL-Gh datasets showed high particle density and side-preferred orientation, the orientation distribution was more biased toward side view in the GroEL-EG than in GroEL-Gh (Fig. 2d,e,g,h,j). However, the resolution of the final map in GroEL-EG (1.99 Å) was higher than those in the others (2.08 and 2.09 Å) (Figs. 2k,l and Supplementary Fig. 4). We also calculated the FSC curves and sphericities of these datasets using the 3DFSC server^16^ and found that the sphericity of GroEL-EG (0.991) was slightly higher than those of GroEL-Gl and GroEL-Gh (0.985 both) (Supplementary Fig. 4). The number fraction of particles used for the final map in those initially picked, which we named the final/initial particle number ratio (FI ratio), should reflect the fraction of protein molecules keeping the intact structures upon blotting and quick freezing in the vitreous thin ice film. In the GroEL-EG dataset, 88.9% (158,485 out of 178,215) of the particles remained after 2D and 3D classifications, but only 19.2% (215,011 out of 1,118,169) and 58.4% (198,677 out of 340,446) did so for the GroEL-Gl and GroEL-Gh dataset, respectively (Fig. 2d and Supplementary Table 1). It should be noted that the number of particles initially picked is strongly influenced by auto-picking parameters. Therefore, our policy was to pick as many particles as possible at the beginning using the same parameters in all three datasets and to classify and discard bad particles later. Many particles were discarded during 2D classification in the GroEL-Gl and GroEL-Gh dataset because the largest number of particles were classified in the blurred 2D class averages (Fig. 2f,g, upper left).

### GroEL particle locations within the ice layer on EG-grid

We collected an electron cryotomography (ECT) dataset from the same GroEL-EG grid to investigate the locations of GroEL particles embedded in the vitreous ice film on the EG-grid (Supplementary Fig. 5). We selected an area of densely packed GroEL particles with some small ice contaminants. It was clear from highly-tilted images that these ice contaminants were located on the surface of vitreous ice. Reconstruction of tomographic images from this dataset revealed three unique layers from the top to bottom. In the top layer, small ice contaminants were observed and no GroEL particles were found. The middle layer 240 Å below the top layer contained densely packed GroEL particles. Unexpectedly, at 225 Å below the middle layer, there were a few GroEL particles that seemed to be bound on the back side of the graphene layer. The estimated distance from the air-water interface to the bottom of the major GroEL layer was ~160 Å, which is reasonable to accommodate a single GroEL layer. These results indicate that GroEL particles were immobilized close to the graphene surface and away from the air-water interface (see also Supplementary Video 2). We also collected an ECT dataset from the GroEL-Gh grid and selected a hole containing both areas with and without graphene support for comparison. The graphene area showed a high density of GroEL particles in a single layer like the EG-grid, demonstrating the compatibility with the EG-grid, while fewer particles were observed in the non-graphene area (Supplementary Fig. 6 and Supplementary Video 3).

### CryoEM image analysis of SARS-CoV-2 spike protein on EG-grid

Next, we examined the performance of the EG-grid on SARS-CoV-2 spike protein, for which the present demand for high-resolution structures is quite high. We prepared three specimen grids: EG-grid (referred to as Spike-EG) and glow discharged graphene grid (referred to as Spike-G), both at a protein concentration of 0.1 mg ml^−1^, and conventional Quantifoil grid (referred to as Spike-Q) at a protein concentration of 0.5 mg ml^−1^. None of the grids were washed with the buffer after protein loading. The particle density was much higher in Spike-EG than that in Spike-Q, although the protein concentration was five times lower for Spike-EG (Fig. 3a,c). The particle density in Spike-G was too low to collect a dataset (Fig. 3b). The 2D class averages showed a wider variety in Spike-EG than those in Spike-Q (Fig. 3d,e), and in agreement with this, Spike-EG showed more uniform orientation distribution than Spike-Q (Fig. 3f,g). The 3D maps reconstructed from these two datasets both showed a conformation of the receptor binding domain (RBD) with 1-up and 2-down (Fig. 2h and Supplementary Fig. 7). Particle images of 150,316 and 58,102 used for the final reconstructions were picked up from 1,163 and 1,029 micrographs, resulting in resolutions of 3.10 Å and 3.23 Å for the Spike-EG and Spike-Q dataset, respectively (Fig. 2i and Supplementary Table 1). The value of NPPI-final for Spike-EG (129.2) was about twice as much as that for Spike-Q (56.5), while the FI ratio values were quite similar (24.9 % and 23.8%) (Supplementary Table 1).

**Fig. 3.**
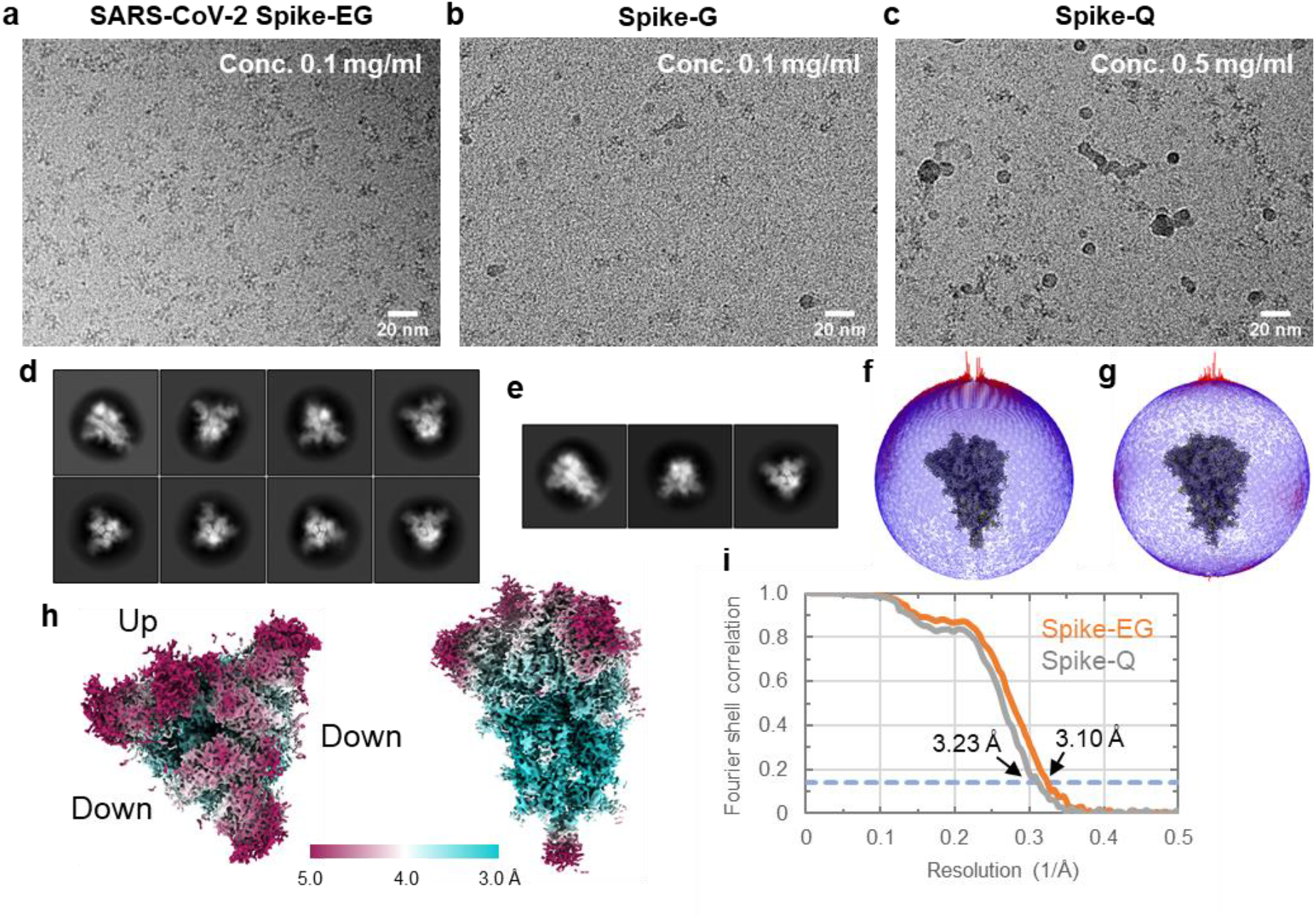
CryoEM image analyses of SARS-CoV-2 spike protein on the EG-grid and Quantifoil grid. **a**–**c**, Typical cryoEM images of spike protein on the EG-grid (spike-EG) (**a**), on the glow discharged graphene grid (spike-G) (**b**), and on the Quantifoil grid (spike-Q) (**c**). **d**,**e**, Selected 2D class averages from the spike-EG dataset (**d**) and the spike-Q dataset (**e**). The class averages are aligned in descending order of particle numbers from left to right and top to bottom. **f**,**g**, Angular distributions of particles used in the final refinement for the spike-EG dataset (**f**) and the spike-Q dataset (**g**). Final 3D maps are also shown for reference. **h**, Two orthogonal views of the final 3D map from the spike-EG dataset: top view of the trimer, left panel; and side view, right panel. The local resolution distribution is colored as in the color bar. **i**, The FSC curves for the final maps of the spike-EG and spike-Q dataset. The dashed blue line indicates the FSC = 0.143 criterion.

### CryoEM analysis of V_1_-ATPase, GAPDH, β-galactosidase, and apoferritin on EG-grid

We applied the EG-grid to the chimeric complex of a soluble domain of V/A-type ATPase (V_1_-ATPase) composed of A_3_B_3_ hexametric ring from *Thermus thermophilus* and DF subunits from *Homo sapiens*, for which the resolution of cryoEM 3D reconstruction has been limited to ~5 Å due to severe preferred orientation (unpublished). We loaded a 1.0 mg ml^−1^ chimeric V_1_-ATPase complex solution onto an EG-grid and made a cryoEM grid by plunge freezing in the same way as we did for GroEL. We saw many protein particles in the micrographs taken at a magnification of 50,000× (Supplementary Fig. 8). A dataset was collected and processed (referred to as V_1_-ATPase-EG). In total, 1,193,032 particles were extracted from 2,052 images (Supplementary Table 1), and 2D class averages showed characteristic hexameric arrangements with a wide variety in particle orientations (Supplementary Fig. 8). During 3D classification, we found two classes, one with and the other without the human DF subunits corresponding to the central rotary shaft. We only selected the class without the shaft, because the particle number of this class was much larger than that of the other. The 3D reconstructed map and model with the shaft will be presented elsewhere. The orientation distribution of the particles used for the final map was much improved (Supplementary Fig. 8) as expected from the 2D class averages. The overall resolution of the final 3D map reconstructed from 238,765 particles reached 3.03 Å (Fig. 4a), and the FI ratio and the NPPI-final were 20.0% and 116.4, respectively (Supplementary Table 1).

**Fig. 4.**
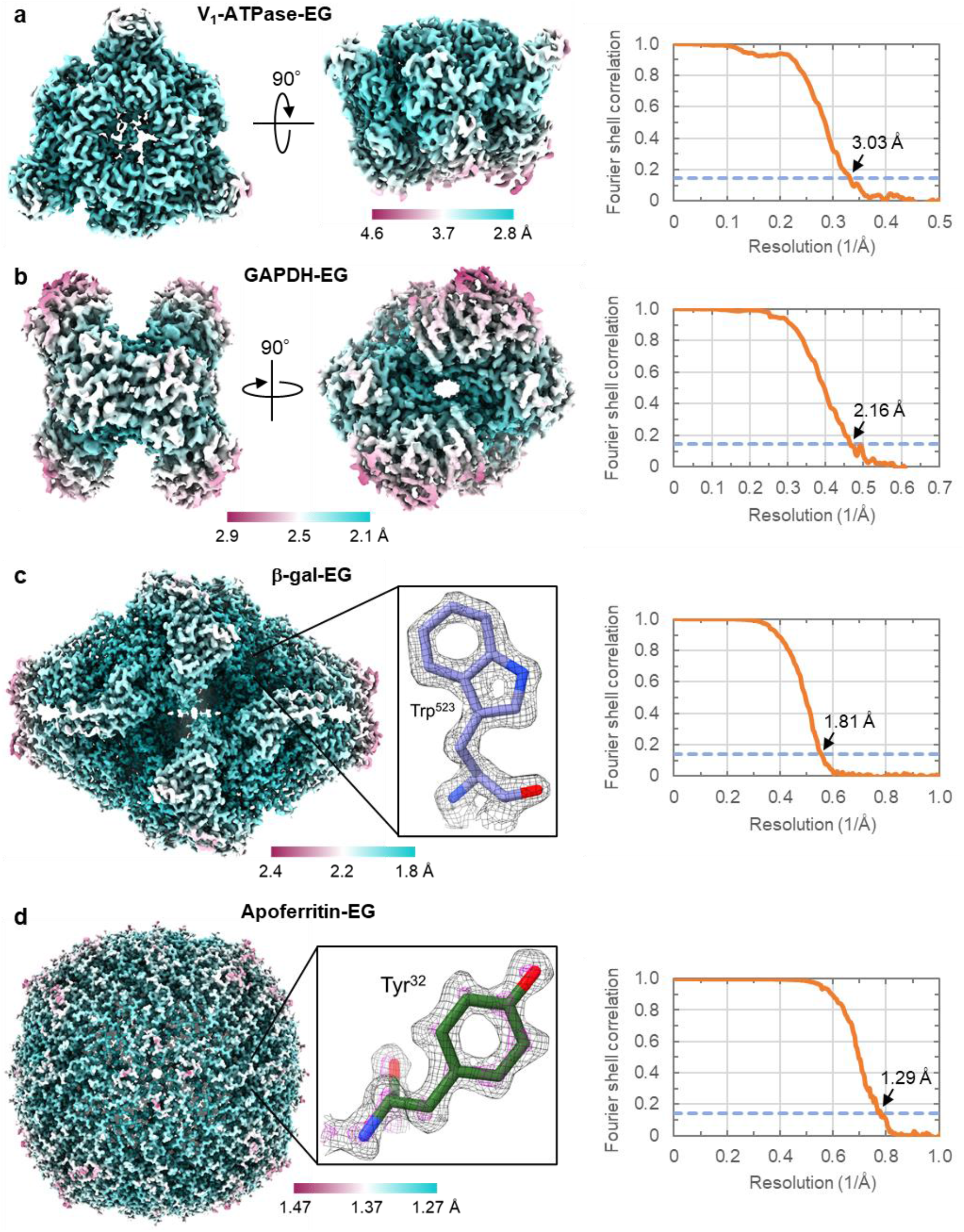
CryoEM analyses of the A_3_B_3_ ring of V_1_-ATPase, GAPDH, β-galactosidase, and apoferritin on EG-grids. **a**–**d**, The final 3D maps from the V_1_-ATPase-EG (**a**), GAPDH-EG (**b**), β-galactosidase-EG (**c**), and apoferritin-EG (**d**) dataset. Two orthogonal views are shown in (**a**) and (**b**). The insets show the density maps and the fitted models of β-galactosidase (PDB entry: 6X1Q) around Trp 523 (**c**) and apoferritin (PDB entry: 6V21) around Tyr 32 (**d**). The map in (**d**) is shown in two contour levels of 0.05 and 0.15 in black and magenta, respectively. The local resolution distributions are colored as in the color bars. Right panels show the FSC curves for the final maps. The dashed blue lines indicate the FSC = 0.143 criterion.

We also applied the EG-grid to the tetramer of glyceraldehyde 3-phosphate dehydrogenase (GAPDH), whose structure has not been solved by cryoEM except for the part of the large ternary complex^31^, probably because of strong preferred orientation with conventional grids (to be published elsewhere). We prepared cryoEM grids by loading an 0.5 mg ml^−1^ GAPDH solution onto an EG-grid and were able to observe densely packed particles in micrographs taken at a magnification of 60,000× (Supplementary Fig. 9). A dataset was collected and processed (referred to as GAPDH-EG). In total, 301,451 particles were extracted from only 241 images (Supplementary Table 1), and 2D class averages showed a wide variation in particle orientations (Supplementary Fig. 9). The orientation distribution of the particles used for the final map was uniform as expected from the 2D class averages. The overall resolution of the final 3D map reconstructed from 88,731 particles reached 2.16 Å (Fig. 4b), and the FI ratio and the NPPI-final were 29.4% and 368.2, respectively (Supplementary Table 1).

We also used the EG-grid to determine the structures of β-galactosidase (β-gal) and apoferritin to validate the efficiency of data collection and capability of reaching high-resolution. We prepared cryoEM grids by loading an aliquot of 1.5 mg ml^−1^ β-gal from *Escherichia coli* or 1.8 mg ml^−1^ mouse apoferritin onto EG-grids, and micrographs taken at a magnification of 100,000× or 120,000× showed a large number of protein particles (Supplementary Figs. 10 and 11). A dataset was collected and processed (referred to as β-gal-EG and apoferritin-EG). In total, 491,997 and 793,398 particles were extracted from 3,242 and 7,500 images for β-gal-EG and apoferritin-EG, respectively (Supplementary Table 1), and the 2D class averages showed a wide variation in particle orientations (Supplementary Figs. 10 and 11). The orientation distributions of the particles used for the final maps were uniform as expected from the 2D class averages. The overall resolution of the final 3D map of β-gal-EG and apoferritin-EG reconstructed from 231,395 and 527,261 particles reached 1.81 and 1.29 Å, respectively (Fig. 4c,d). The central region of the β-gal-EG density map had a sufficiently high resolution and quality to show the holes in the center of two aromatic rings of Trp residues (Fig. 4c, inset). The density map of apoferritin-EG also had a sufficient high resolution and quality to display individual atoms as blobs at a relatively high contour level, as indicated by the contour in magenta for a Tyr residue (Fig. 4d, inset). The FI ratio and the NPPI-final of β-gal-EG and apoferritin-EG were 47.0% and 66.5%, and 71.4 and 70.3, respectively (Supplementary Table 1). The overall resolution of β-gal-EG is almost the same as the highest resolution 1.82 Å of the map deposited in the EMDB to date, which was reconstructed from 257,202 particles extracted from 4,949 images^32^. So the EG-grid allowed to reach the highest resolution of β-galactosidase with 40% less number of micrographs.

## Discussion

Oxidation of graphene films on EM grids without damaging is not an easy task. In fact, the plasma method (glow discharge) has been reported to cause graphene to collapse in a short time^14^. It has also been reported that UV/O_3_ or KMnO_4_ treatment leads to the introduction of carboxy groups, implying the cleavage of C=C bonds in graphene, and that was confirmed by the formation of nanopores^13^. As can be seen in these reports, it is difficult to oxidize chemically stable graphene to introduce oxygen functional groups on the surface without disruption of its atomic layer sheet structure. In our method, however, C-O single bond groups (hydroxy groups) are more preferentially introduced compared to C=O type functional groups such as carboxy groups (Supplementary Fig. 1), and therefore the intrinsic sheet structure of graphene is better maintained. We also confirmed that these hydroxyl groups react with ECH, which further reacts with amine compounds (Supplementary Fig. 2).

As we tested the EG-grid for many samples, we found that the effectiveness of the EG-grid somewhat varied from sample to sample. For GroEL, both GroEL-EG and GroEL-Gh showed densely packed particles and large values for NPPI-final, which means that many particles are kept attached even after buffer washing in some area of graphene surface without epoxidation. However, when a much lower concentration GroEL solution than the above experiment was applied, many of the GroEL particles were washed away by buffer washing even with the EG-grid. Therefore, despite the demonstration of covalent bonding between a small molecule with an amino group and the epoxy group introduced to the graphene surface by ECH, immobilization of protein particles on the EG-grid by the covalent bonding of their amino groups to the epoxy groups appeared to be not so efficient at least for GroEL. In addition, the orientation distribution of GroEL on a cryoEM grid was strongly affected by the particle density rather than the epoxidation of graphene. The biggest difference between the EG-grid and hydrophilized graphene grid was the value of FI ratio. We suspect that the much higher FI ratio and the better resolution of the final 3D map (1.99 Å) of GroEL-EG reflects the intactness of protein particles kept away from the graphene surface by the linker introduced during epoxidation. Another example demonstrating the effectiveness of the EG-grid is SARS-CoV-2 spike protein. The Spike-EG images showed much higher particle density at a very low concentration (0.1 mg ml^−1^) compared to those of Spike-G and Spike-Q and an improved orientation distribution of the particles. Although the overall resolution of the final 3D map was not dramatically improved by using the EG-grid mainly because of the flexible nature of the RBD in the spike protein, the EG-grid was still beneficial in terms of saving the amount of sample and the time for grid screening and data collection.

In other examples, V_1_-ATPase-EG and GAPDH-EG represented an improved orientation distribution of the particles. GAPDH-EG also proved that near 2 Å resolution is reachable in ~150 kDa protein only from ~240 images, which we can collect in about half an hour now. β-gal-EG and apoferritin-EG demonstrated the possibility of high resolution structure determination using the EG-grid. Of course, there may be some cases where the EG-grid is not effective enough to reach high resolution. For example, we applied the EG-grid to hemagglutinin (HA), which suffered from preferred orientation and needed grid tilting data collection for 3D reconstruction^16^. HA on the EG-grid showed a large number of particles on each micrograph and 2D class averages with an improved orientation distribution (many side views, Supplementary Fig. 12), but the resolution of the 3D map was limited to 6–7 Å.

In summary, the parameters indicating the superiority of the EG-grid seem to vary from protein to protein, but the clear and general advantage of the EG-grid is that much more protein particles can be observed in the same size of viewing area with a more uniform orientation distribution than conventionally used holey carbon grids, and in most cases, one or two grid preparation is sufficient, greatly reducing the time required for grid screening for cryoEM data collection for high-resolution 3D reconstruction. The difference in the particle density would be due to the difference in the affinity between proteins and the epoxidized graphene surface. Since proteins are likely to be adsorbed on the EG-grid through multiple contact points, the shape, flatness, charge, and stability of protein may affect the affinity, altering the particle density on the grid as well as their orientation distribution. We need more examples, experiences, and further investigation on this point. But in many cases, the method of immobilizing proteins on the EG-grid prepared by ClO_2_^•^ oxidation is expected to dramatically improve the efficiency and resolution of structure determination by cryoEM, especially for those many proteins suffering from preferred orientation and/or low yield. Moreover, the ClO_2_^•^ oxidized graphene surface can be chemically modified easily in many different ways to develop different types of grids with different surface functionalities that would further expand the potential of cryoEM towards much wider applications, for example, by allowing easier and quicker on-the-grid sample purification.

## Methods

### Graphene grid fabrication

We prepared graphene grids using Quantifoil grids based on the protocol described previously with some modifications^12^. PMMA-coated CVD graphene, either 2, 4 or 6-layers, on a Cu foil (AirMembrane), or PMMA-coated single-layer CVD graphene on a Cu foil from any suppliers (ACS Material, Graphenea, SIGMA-Aldrich etc.) can be used. If necessary, the back-side (the opposite side of PMMA) of graphene was removed by mechanical polishing using a waterproof abrasive paper (P3000, NIHON KENSHI Co., Ltd.). The Cu foil was etched by floating the PMMA-coated CVD graphene on a 0.5 M ammonium persulfate (APS) solution for more than 30 min. Then, the PMMA-coated graphene was scooped up on a filter paper, transferred to water and left for 10 min. This washing process was repeated twice. Or, a much easier and quicker method is to use Trivial Transfer Graphene™ (ACS Material), which can be transferred to water directly without polishing, etching, and washing. The graphene was transferred to water in a stainless container in which a number of Quantifoil grids (R1.2/1.3 Au 200 mesh) were aligned at the bottom with their carbon side facing up. The water was slowly drained, and the graphene layer was deposited on the grids. After air-dried for more than 1 hour, each grid was separated and baked at 130 °C for 15 min. The grids were immersed in acetic acid twice for PMMA removal: for 2–3 hours in the first step and then overnight. Then the grids were soaked in isopropanol for 10 min, air-dried on a filter paper, and baked at 100 °C for 10 min. The grids were stored in a grid box in a desiccator.

### Oxidation of graphene grid

The graphene grids prepared as above were oxidized by photo activated ClO_2_^•^ gas generated from an aqueous solution (20 ml) of NaClO_2_ (200 mg) and 37% HCl (aq) (100 μl) by irradiation with an UV LED lamp (λ = 365 nm, 20 mW/cm^2^) for 10 min. A square-shaped dish-like glassware with an inner and outer compartment that we developed before (shown in Fig. 6 of ^33^) was used to perform this oxidation process. The aqueous solution of NaClO_2_ was filled into the outer compartment, the graphene grids were laid in the inner compartment, and a glass plate was put on the glassware to seal the entire chamber. To avoid any damage on the graphene layer by direct photoirradiation, an aluminum foil was laid on the grids to shield them from the UV light (see also Supplementary Video 1). The total reaction volume can be reduced to 5 ml. The graphene on a silicon wafer were also oxidized in the same way. Because the ClO_2_^•^ gas is toxic and potentially explosive especially at high concentration, all the oxidation process should be performed under controlled concentration in a fume hood. Plasma treated hydrophilized graphene on a silicon wafer were also prepared as a control for XPS measurement by glow discharge (30 mA, 20 s) using a JEC-3000FC sputter coater (JEOL) with an aluminum foil mask to shield them from plasma irradiation.

### Fluorination of graphene grid

The preparation of EG-grid (surface functionalization of oxidized graphene by 1% ECH) was performed as described in manuscript for negative staining. A 3 μl aqueous solution of 1% 1H,1H-undecafluorohexylamine (UFHA) was loaded on the EG-grid. After 5 min, the solution was blotted with a filter paper and the grid was washed with 5 μl water for 3 times. After washing process, the fluorinated graphene grid was dried in vacuo overnight.

### Surface analysis of graphene

Raman spectra were obtained with an NRS-3100T laser Raman spectrometer (JASCO) with a 532 nm laser at room temperature. Fourier-transform infrared (FTIR) spectroscopic measurements were performed using a Spectrum Two spectrometer (Perkin-Elmer) equipped with an UATR two attachment. All the spectra were acquired at a resolution of 4 cm^-1^ over 16 scans in a scan range of 600–4000 cm^-1^. The surface chemical composition of the graphene was determined using a JPS-9010MC XPS instrument (JEOL). The XPS insert parameters included the power of analysis (wide: 75 W, narrow: 150 W) and monochromatic Mg Kα radiation. The survey and high-resolution XPS profiles were obtained at the constant analyzer pass energies of 160 and 10 eV, respectively. The CasaXPS Version 2.3.15 software was used for the peak-differentiation-imitating analysis of the C1s narrow XPS profile. Two independent experiments were conducted to obtain the standard deviation of the data. The static water contact angles of the graphene surfaces were determined using a Drop Master DM300 (Kyowa Interface Science) contact angle meter. A 1.0 μl of water droplet was placed on the surface of the graphene on the grid, and the contact angle was determined 5 s (scanning time) after the attachment of the droplet.

### Protein preparation

GroEL from *Escherichia coli* was purchased from TaKaRa Bio Inc. (Product code 7330). A 5 mg powder sample of GroEL was dissolved with 500 μl of GroEL buffer (25 mM HEPES pH 8.0, 50 mM NaCl), and it was purified using a Superdex 200 Increase 10/300 GL gel filtration column with the GroEL buffer. The main peak was collected and concentrated to 3.2 mg ml^−1^, flash-frozen with liquid nitrogen and stored at −80°C.

The codon-optimized gene of SARS-CoV-2 Spike (S) protein (GenBank: QHD43416) was designed for expression in mammalian cells and synthesized from GeneArt DNA Synthesis (Thermo). For expression of recombinant S protein, the sequence encoding the S ectodomain (residues 1–1208) with proline substitutions at residues 986 and 987, a “GSAS” substitution at furin cleavage site (residues 682–685), and C-terminal foldon trimerization motif followed by an octa-histidine tag was cloned into a pcDNA3.1 expression vector (Invitrogen). Further, S protein D614G mutation was introduced by inverse PCR method. Recombinant S proteins were transiently expressed in Expi293f cells (Thermo) maintained in HE400AZ medium (Gmep, Japan). The expression vector was transfected by using Gxpress 293 Transfection Kit (Gmep, Japan) as following the manufacturer’s protocol. After 5 days post-transfection, culture supernatants were harvested, and His-tagged S proteins were purified by Ni^2+^ affinity chromatography using Ni Sepharose 6 Fast Flow (Cytiva), followed by size exclusion chromatography using Superdex 200 Increase 10/300 GL (Cytiva) equilibrated with a buffer containing 50 mM HEPES (pH 7.0) and 200 mM NaCl.

*Escherichia coli* strain BL21-CodonPlus-RP (Stratagene) was used for expression of the human DF-V_1_-ATPase chimeric complex. The recombinant complex was isolated as described previously^34,35^. The expression plasmids for V_1_-ATPase containing DF from *Homo sapiens* were constructed by the same method as described previously^34^. The genes encoding the D and F subunits were amplified from human cDNA. The amplified fragment was then digested with appropriate restriction enzymes and inserted into the corresponding region of the *Thermus thermophilus* V_1_-ATPase expression plasmid^35^. The mutant V_1_-ATPase (A-His8/ΔCys, A-C255A/A-S232A/A-T235S, FS54C) was used for preparation of the complex.

The gene of human GAPDH was cloned into pET-28a(+) expression vector (Novagen) with an N-terminal His-tag followed by the Tobacco Etch Virus (TEV) protease cleavage site. The recombinant protein was expressed using an *Escherichia coli* strain, BL21 Star™ (DE3) (Thermo Fisher Scientific). The transformed cells were cultured in Luria-Bertani (LB) medium with 50 μg ml^−1^ Kanamycin at 37 °C until the OD_600_ reached ~0.6. After adding 1 mM isopropyl-β-D-thiogalactopyranoside (IPTG) and culturing for 16 h at 37 °C, cells were harvested by centrifugation, flash-frozen with liquid nitrogen, and stored at −80 °C. Frozen bacteria pellets were resuspended in a buffer containing 20 mM Tris-HCl pH 8.0, 30 mM NaCl, 1 mM 1,4-dithiothreitol (DTT) supplemented with 10 μl Benzonase, 1 mg/ml lysozyme, and EDTA-free cOmplete™ protease inhibitor cocktail (Roche) and lysed by sonication. The suspension was centrifuged for 30 min at 200,000 × *g* and 4 °C. The supernatant was collected and purified using a HisTrap HP 5 ml column (GE Healthcare) equilibrated with a buffer containing 20 mM Na_4_P_2_O_7_ pH 7.4, 500 mM NaCl, 30 mM imidazole, and subsequently eluted with the same buffer containing 500 mM imidazole. The desired fractions were collected and dialyzed overnight at 4 °C against a buffer composed of 20 mM Na_4_P_2_O_7_ pH 7.4, 200 mM NaCl supplemented with TEV protease. The protein was further purified using a HiLoad 16/600 Superdex 200 column (GE Healthcare) equilibrated with 20 mM Tris-HCl pH 8.0, 30 mM NaCl, 1 mM Tris(2-carboxyethyl)phosphine (TCEP). The desired fractions were collected, concentrated to ~10 mg ml^−1^, flash-frozen with liquid nitrogen, and stored at −80 °C.

β-galactosidase (β-gal) was purchased from SIGMA-Aldrich (Product code G5635) and dissolved with β-gal buffer (25 mM HEPES pH 8.0, 50 mM NaCl, 2 mM MgCl_2_, 1 mM TCEP) to a final concentration of 4.0 mg ml^−1^. IPTG was supplemented to a final concentration of 5 mM, and the solution was incubated on ice for 30 min. β-gal was purified using Superdex 200 Increase 10/300 GL column (GE Healthcare) with the same buffer. The main peak was collected and concentrated to 2.0 mg ml^−1^, flash-frozen with liquid nitrogen and stored at −80 °C.

Mouse apoferritin was expressed using mFth1-pET24a plasmid and an *Escherichia coli* strain, BL21 Star™ (DE3) (Thermo Fisher Scientific). After the plasmid was transformed, the cells were cultured in 4 ml of LB medium with 50 μg ml^−1^ Kanamycin and 0.5% glucose overnight. The culture was transferred into 500 ml of fresh culture medium and cultured at 37°C until the optical density at 600 nm reached 0.5. Then, 1 mM IPTG was added, and the cultivation was continued for 4 hours at 37 °C. Cells were harvested and resuspended in buffer A (30 mM HEPES pH 7.5, 300 mM NaCl, 1 mM MgSO_4_) supplemented with 1 mg ml^−1^ lysozyme and lysed by sonication. The suspension was centrifuged, and the supernatant was incubated at 70 °C for 10 min. After centrifugation and collecting supernatant again, the protein was precipitated by adding ammonium sulfate to a final concentration of ~53% and stirring on ice for 30 min. After centrifugation and collecting pellet, the pellet was washed with buffer A supplemented with ~53% ammonium sulfate. This washing step was repeated twice. The washed pellet was resuspended in 2 ml of phosphate buffered saline (PBS) and dialyzed against 500 ml of PBS at 4°C overnight. Precipitation was removed by ultracentrifugation, and the solution was concentrated to ~0.9 ml, which was further purified using a Superose 6 10/300 column (GE Healthcare) equilibrated with 20 mM HEPES pH 7.5, 150 mM NaCl. The desired fractions were collected separately, flash-frozen with liquid nitrogen and stored at −80°C.

Influenza hemagglutinin trimer (HA) (H3N2, A/Hong Kong/1/1968) was purchased from MyBioSource (catalog number: MBS434205) and dissolved in 1x PBS buffer to a concentration of 0.5 mg ml^−1^.

### Epoxidation of oxidized graphene grid and negative staining

A 3 μl solution of 1% (v/v) epichlorohydrin (ECH) in water was loaded on the graphene side of an oxidized graphene grid held with a tweezer. After 5 min, the ECH solution was blotted with a filter paper, and the grid was washed with 5 μl of ultrapure water (or protein buffer not containing a compound with amino group such as Tris) three times. Then, 3 μl of GroEL solution (0.3 mg ml^−1^) was loaded on the graphene side of the grid. After 5 min, the protein solution was blotted, and the grid was washed with 5 μl of the same buffer three times (see also Supplementary Video 1). Then the grid was immediately stained with 3 μl of 2% uranyl acetate solution. Images were taken using a JEM-1011 electron microscope (JEOL) at 100 kV. It is preferable to perform the epoxidation process just before the protein loading step, because the epoxidized graphene is no longer usable after a few days.

### CryoEM grid preparation

The surface functionalization of oxidized graphene by 1% ECH and the protein loading step were performed as described above for negative staining. Protein concentration was 1.5 mg ml^−1^ for the GroEL-EG and the β-gal-EG, 0.1 mg ml^−1^ for the SARS-CoV-2 Spike-EG, 1.0 mg ml^−1^ for the V_1_-ATPase-EG, 0.5 mg ml^−1^ for the GAPDH-EG and the HA, and 1.8 mg ml^−1^ for the apoferritin-EG. For SARS-CoV-2 spike protein, the stock sample was diluted with the buffer containing 20 mM HEPES (pH 7.4) and 150 mM NaCl. In the washing step after protein loading, the final (third) washing buffer on the grid was not blotted, and the tweezer holding the grid was placed in a chamber of Vitrobot Mark IV (Thermo Fisher Scientific) equilibrated at 4°C and 100% humidity. For the Spike-EG, GAPDH-EG, and HA, the grid was not washed after the protein loading. After a liquid ethane container was placed, the grid was blotted for 1.5 s (for all samples other than the Spike-EG) or 2.0 s (for the Spike-EG) and immediately plunged into liquid ethane. Excessive ethane was manually blotted with a filter paper, and the grid was stored in liquid nitrogen. For comparison, non-oxidized graphene grid (for the GroEL-Gl, GroEL-Gh, and Spike-G) or Quantifoil grids (R1.2/1.3 Cu 200 mesh, for the Spike-Q) was hydrophilized by glow discharge using a JEC-3000FC sputter coater (JEOL). To avoid graphene breakage, a loose mask made of a thin aluminum foil was placed above the grids during the glow discharge for the graphene grid. A tweezer holding the grid was placed in a chamber of Vitrobot and after the liquid ethane container was placed, 3 μl of 1.5 mg ml^−1^ (for the GroEL-Gl) or 1.0 mg ml^−1^ (for the GroEL-Gh) GroEL, or 0.1 mg ml^−1^ (for the Spike-G) or 0.5 mg ml^−1^ (for the Spike-Q) spike protein solution was loaded on the graphene side. Then the grid was blotted for 1.5 s (for the GroEL-Gl and GroEL-Gh) or 2.0 s (for the Spike-G and Spike-Q) and immediately plunged into liquid ethane. Excessive ethane was manually blotted with a filter paper, and the grid was stored in liquid nitrogen.

### CryoEM data collection for SPA

All the cryoEM image datasets were acquired using SerialEM^36^ and JEM-Z300FSC (CRYO ARM 300, JEOL) operated at 300 kV with a K3 direct electron detector (Gatan, Inc.) in the CDS mode. The Ω-type in-column energy filter was operated with a slit width of 20 eV for zero-loss imaging. For the GroEL-EG, GroEL-Gl, GroEL-Gh, and GAPDH-EG dataset, nominal magnification was 60,000×. Defocus varied between −0.5 and −2.0 μm. Each movie was fractionated to 40 frames (0.07 s each, total exposure 2.8 s) with a total dose of 40 e^−^/Å^2^. For the Spike-EG, Spike-Q, and HA dataset, nominal magnification was 60,000×. Defocus varied between −0.5 and −2.0 μm. Each movie was fractionated to 60 frames (0.05 s each, total exposure 3.0 s) with a total dose of 60 e^−^/Å^2^. For the V_1_-ATPase-EG dataset, nominal magnification was 50,000×. Defocus varied between −1.0 and −2.5 μm. Each movie was fractionated to 48 frames (0.084 s each, total exposure 4.0 s) with a total dose of 62 e^−^/Å^2^. For the β-gal-EG dataset, nominal magnification was 100,000×. Defocus varied between −0.5 and −2.0 μm. Each movie was fractionated to 40 frames (0.065 s each, total exposure 2.6 s) with a total dose of 40 e^−^/Å^2^. For the apoferritin-EG dataset, nominal magnification was 120,000×. Defocus varied between −0.3 and −1.3 μm. Each movie was fractionated to 64 frames (0.025 s each, total exposure 1.6 s) with a total dose of 40 e^−^/Å^2^. More parameters are summarized in Supplementary Table 1.

### SPA image processing

The movies were motion corrected using MotionCor2^37^, and CTFs were estimated using CTFFIND 4.1^38^. All the following steps were performed using RELION 3.1^39^. Only the images whose CTF max resolutions were beyond 5 Å were selected, and the resulting number of images are referred to as Initial images. Laplacian of Gaussian (LoG)-based auto-picking was performed for small number of images to generate the 2D class averages, which were then used as a template for second auto-picking for all images. Picked particles were extracted with 1–4 x binning, and the resulting number of particles was referred to as the initial particle number. Several rounds of 2D classification into 100 classes were carried out to remove bad particles. For V_1_-ATPase-EG, 2D classification into 50 classes was repeated three times. Initial 3D model was generated with C1 symmetry, and then D7 (for GroEL) or D2 (for GAPDH and β-gal) symmetry was applied to the resulting maps. No symmetry was applied for SARS-CoV-2 spike protein and V_1_-ATPase. 3D classification (into 4 classes) with C1 symmetry was conducted to exclude bad particles, and the number of remaining particles was referred to as the final particle number except the GAPDH-EG and β-gal-EG datasets. The remaining particles were re-extracted with a larger box size, and 3D auto-refine applying D7 (for GroEL) or D2 (for β-gal) symmetry was performed. After a soft mask around the map was generated, the resulting map was post-processed and sharpened. After several rounds of CTF refinement and Bayesian polishing with divided optics groups of 8 (for GAPDH-EG), 10 (for GroEL-EG and Spike-Q), 11 (for GroEL-Gl and V_1_-ATPase-EG), 12 (for Spike-EG), 13 (for GroEL-Gh), and 16 (for β-gal-EG), 3D auto-refine and post-processing were conducted again. For the GAPDH-EG and β-gal-EG datasets, particles were re-selected with rlnMaxValueProbDistribution=0.05 criteria (the number of remaining particles was referred to as the final particle number), and 3D auto-refine and post-processing were conducted to generate the final map. The map resolution was estimated by the gold-standard Fourier shell correlation (FSC = 0.143) criterion. The FSC curves and sphericities were calculated using 3DFSC server^16^ (https://3dfsc.salk.edu).

For apoferritin, in total 793,398 particles picked by using 2D class averages template were extracted from 7,500 images (Supplementary Table 1) with a box of 512 x 512 pixels and divided into 25 groups as optical groups according to the time of data collection. After the particle selection by 2D classification, an initial model was generated, and the selected particles were subjected to several rounds of 3D auto-refine with O symmetry and post-processing with CTF refinement and Bayesian polishing. Then the particles were regrouped into 12 by the range of beam tilt X, Y (0.1 milli-radian increments) and further regrouped into 300 (= 12 * 25: 5 x 5 multi-hole shots) by image shift. Regrouped particles were subjected to another round of CTF refinement, 3D auto-refine, and post-processing. Then the particles were further selected with rlnMaxValueProbDistribution = 0.04 criteria, and the selected 527,261 particles were subjected to another rounds of Bayesian polishing with a larger box size (600 x 600), 3D auto-refine, and post-processing. Finally, the Ewald sphere correction and another round of post-processing was applied to generate the final map. Structural analysis and figure preparation were performed using UCSF Chimera^40^ and ChimeraX^41^.

### Tomography

The datasets for tomography were acquired for the same grid as the GroEL-EG or GroEL-Gh dataset was collected using JEM-Z300FSC (CRYO ARM™ 300, JEOL) operated at 300 kV with a K3 direct electron detector (Gatan, Inc.) in CDS mode. The Ω-type in-column energy filter was operated with a slit width of 20 eV for zero-loss imaging. For GroEL-EG, the nominal magnification was 40,000×, corresponding to 1.25 Å/pixel. Defocus was set to −3.0 μm. The tilt angle varied from −60° to 60° with 3° increment to acquire 41 movies using SerialEM^36^, and the order of tilt series was as the following tilt angle in degree: (0, 3, 6, 9) -> (−3, −6, −9) -> (12, 15, 18) -> (−12, −15, −18) -> (21, 24, 27) -> (−21, −24, −27) -> (30, 33, 36) -> (−30, −33, −36) -> (39, 42, 45) -> (−39, −42, −45, −48, −51, −54, −57, −60) -> (48, 51, 54, 57, 60). Each movie was fractionated to 5 frames (0.1 s each, total exposure 0.5 s per tilt angle) with a total dose of 98 e^−^/Å^2^. For GroEL-Gh, the nominal magnification was 20,000 ×, corresponding to 2.37 Å/pixel. Defocus was set to −3.0 μm. The tilt angle varied from −60° to 60° with 3° increment to acquire 41 movies using SerialEM^36^, and the order of tilt series was as the following tilt angle in degree: (0, 3, 6) -> (−3, −6) -> (9, 12) -> (−9, −12) -> (15, 18) -> (−15, −18) -> (21, 24) -> (−21, −24) -> (27, 30) -> (−27, −30) -> (33, 36) -> (−33, −36) -> (39, 42) -> (−39, −42) -> (45, 48, 51, 54, 57, 60) -> (−45, −48, −51, −54, −57, −60). Each movie was fractionated to 5 frames (0.25 s each, total exposure 1.25 s per tilt angle) with a total dose of 40 e^−^/Å^2^. Tomograms were reconstructed using IMOD^42^. Images were aligned without using gold nanoparticles by patch tracking (for GroEL-EG) or by tracking ice contaminations as seeds (for GroEL-Gh).

## Data availability

Density maps are available at EMDB with accession codes EMD-31310 (GroEL-EG), EMD-31311 (GroEL-Gl), EMD-32159 (GroEL-Gh), EMD-32160 (SARS-CoV-2 Spike-EG), EMD-32161 (SARS-CoV-2 Spike-Q), EMD-31312 (V_1_-ATPase-EG), EMD-32162 (GAPDH-EG), EMD-31313 (β-galactosidase-EG), and EMD-31314 (apoferritin-EG). Other data supporting this study are available from the corresponding authors on reasonable request.

## Acknowledgments

We thank Haruaki Yanagisawa and Masahide Kikkawa for the plasmid of mouse apoferritin, Miki Kinoshita for expression and purification of apoferritin, Chun-ming Zhang, Yusuke Yasuda, and Ryo Kanno for expression, purification, and data collection of GAPDH, Kanato Arita for video preparation, Masataka Hasegawa and Yoshinori Koga for CVD graphene sheet samples produced by AirMembrane, and JEOL engineers and staffs for their help and efforts in the development of the CRYO ARM electron cryomicroscope for high-throughput data collection. This work was supported by: JST OPERA (Open Innovation with Enterprises, Research Institute and Academia) grant JPMJOP1861 (T.I., K.N.); NEDO (New Energy and Industrial Technology Development Organization) grant 16102003-0 (T.I.) and 17101509-0 (H.A.); JSPS KAKENHI grant JP25000013 (K.N.) and JP20K22630 (J.F.); AMED BINDS (Platform Project for Supporting Drug Discovery and Life Science Research (BINDS) grant JP19am0101117 (K.N.); AMED CiCLE (Cyclic Innovation for Clinical Empowerment) grant JP17pc0101020 (K.N.); JEOL YOKOGUSHI Research Alliance Laboratories of Osaka University (K.N.).

## Author contributions

T.I. and K.N. conceptualized, administrated, and supervised the project. J.F., F.M., H.A., and M.M. designed the experiments. J.F., H.A. M.M. and S.K. prepared and analyzed EG-grid. J.F., S.K., I.A., and J.K. prepared the protein samples. J.F., F.M., J.K. and T.K. performed cryoEM experiments. J.F., F.M. and H.A. prepared the graphics and illustrations. J.F., F.M., H.A., T.I. and K.N. wrote the first draft of the manuscript. J.F., F.M., H.A., M.M., I.A., J.K., Y.M., T.K., K.N., and T.I. reviewed and commented on the manuscript. All authors approved the reviewed manuscript.

