## Supplementary Information for "Epoxidized graphene grid for high-throughput high-resolution cryoEM structural analysis"

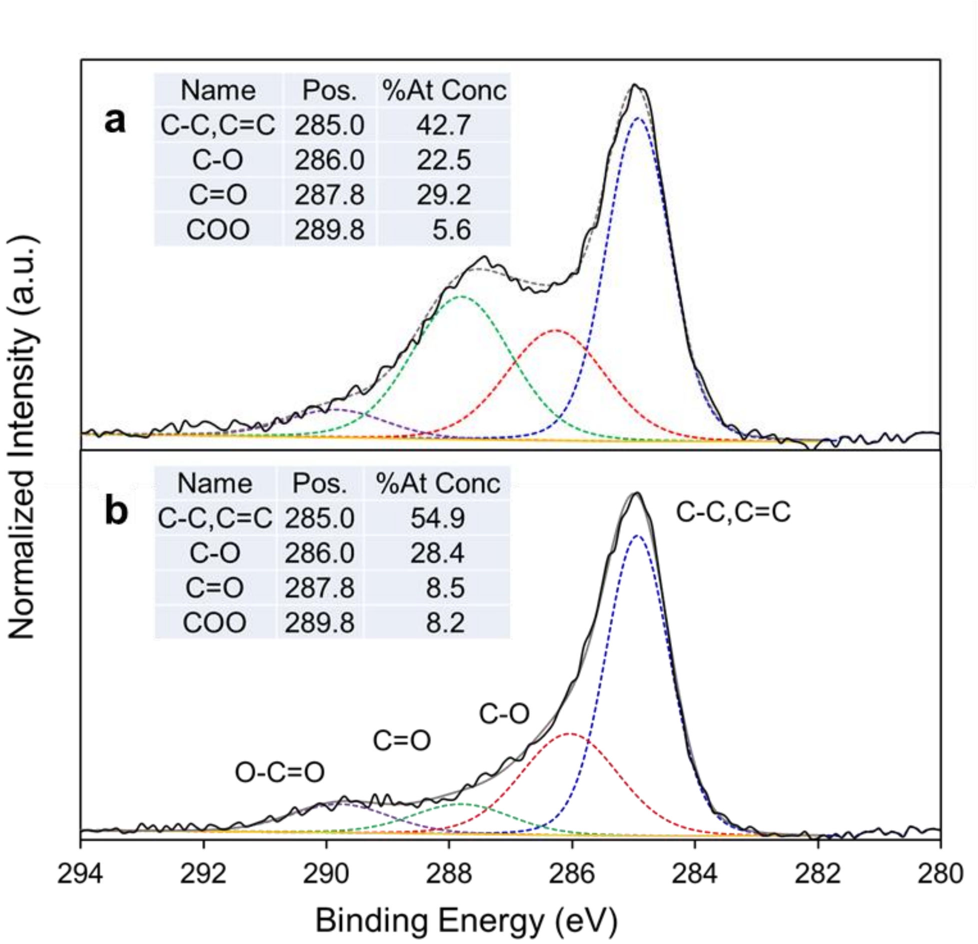


**Supplementary Figure 1. XPS spectra of chemically modified graphene. a**,**b**, C 1s high-resolution spectra of plasma-treated (**a**) and $ClO_2^{\bullet}$-treated (**b**) graphene on a silicon wafer. The inset shows binding energies of the C 1s electrons in the functional groups that were used in the fitting procedure and the composition ratios.

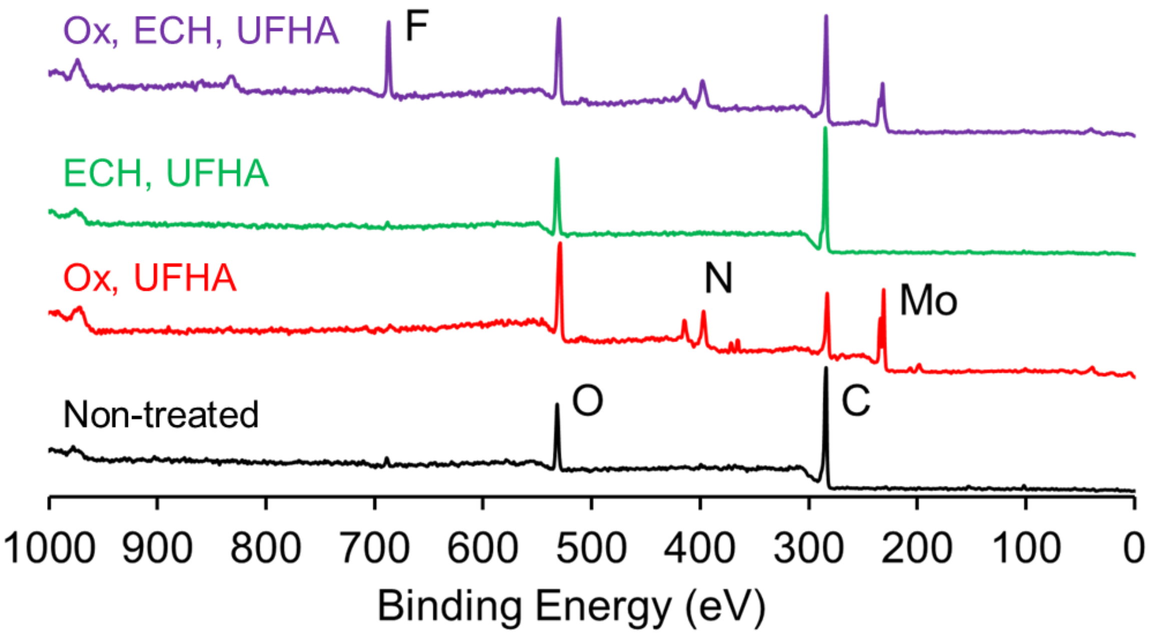


**Supplementary Figure 2. XPS spectra of graphene grids.** XPS spectra of graphene (black line), UFHA-treated oxidized graphene (red line), epichlorohydrin- and UFHA-treated graphene (green line), on the Quantifoil Mo grid, and UFHA-treated EG-grid (purple line).

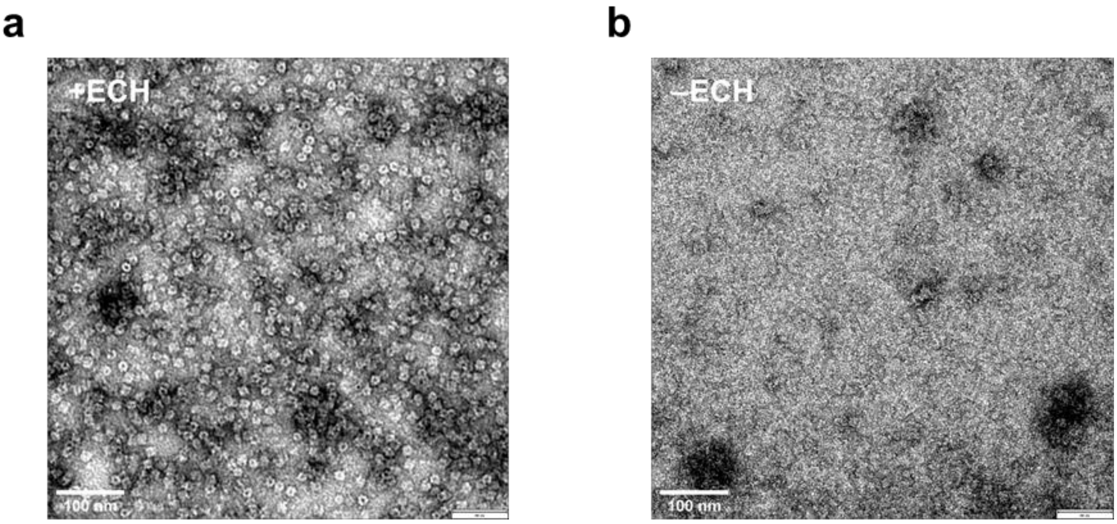


**Supplementary Figure 3. Protein binding activity of the EG-grid. a**,**b**, Typical electron micrographs (30,000 x) of GroEL negatively stained (**a**) on an EG-grid after washing by GroEL buffer three times before staining and (**b**) on a grid prepared in the same way as (**a**) except omitting the epoxidation process by ECH.

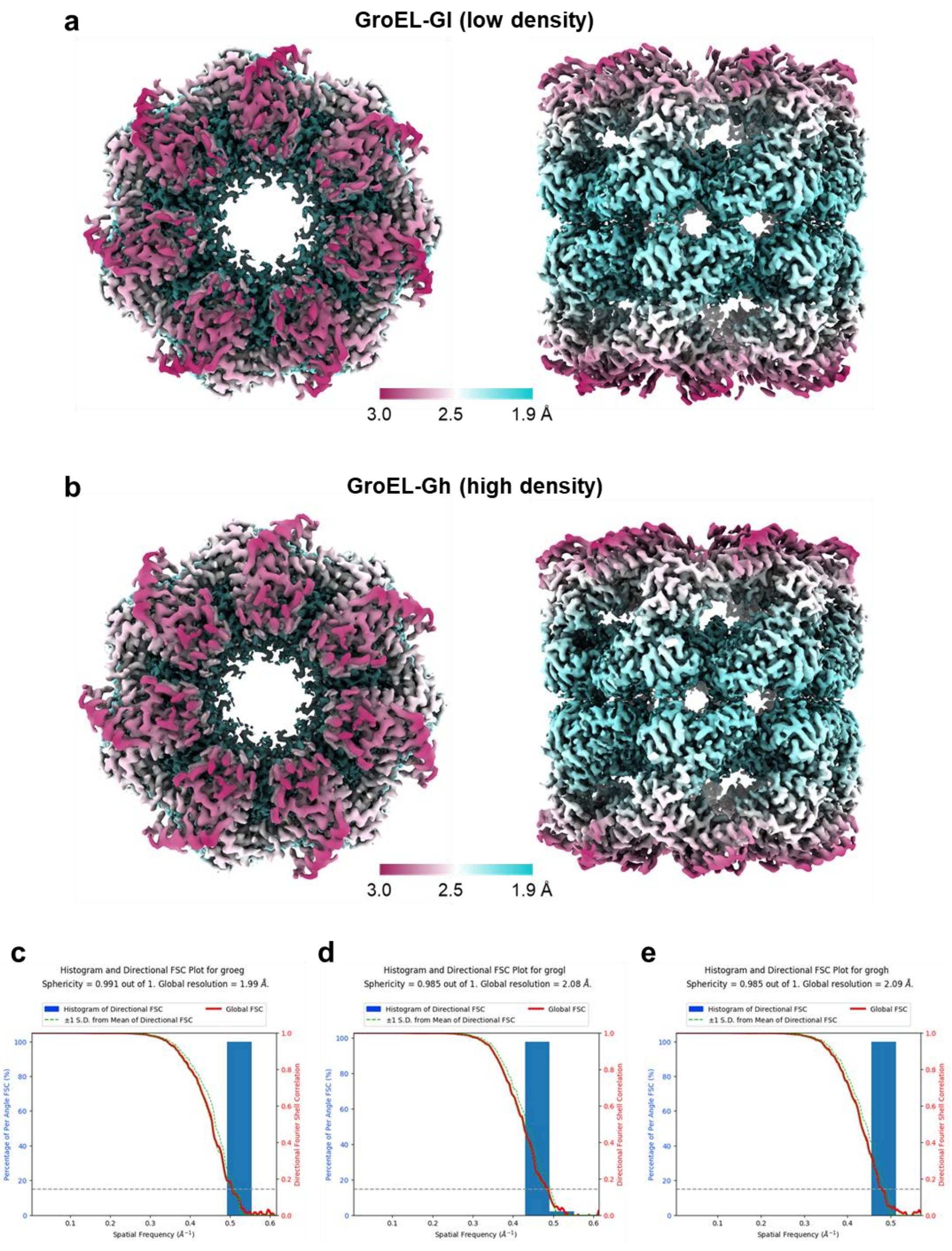


**Supplementary Figure 4. CryoEM image analysis of the GroEL. a**,**b**, Final 3D maps of GroEL-Gl (**a**) and GroEL-Gh (**b**) dataset in two orthogonal views: top view, left panels; and side view, right panels. The local resolution distributions are colored as in the color bars. **c**-**e**, FSC curves and sphericities calculated by the 3DFSC server (https://3dfsc.salk.edu) for the final maps of the GroEL-EG (**c**), GroEL-Gl (**d**), and GroEL-Gh (**e**) datasets. The dashed line indicates the FSC = 0.143 criterion.

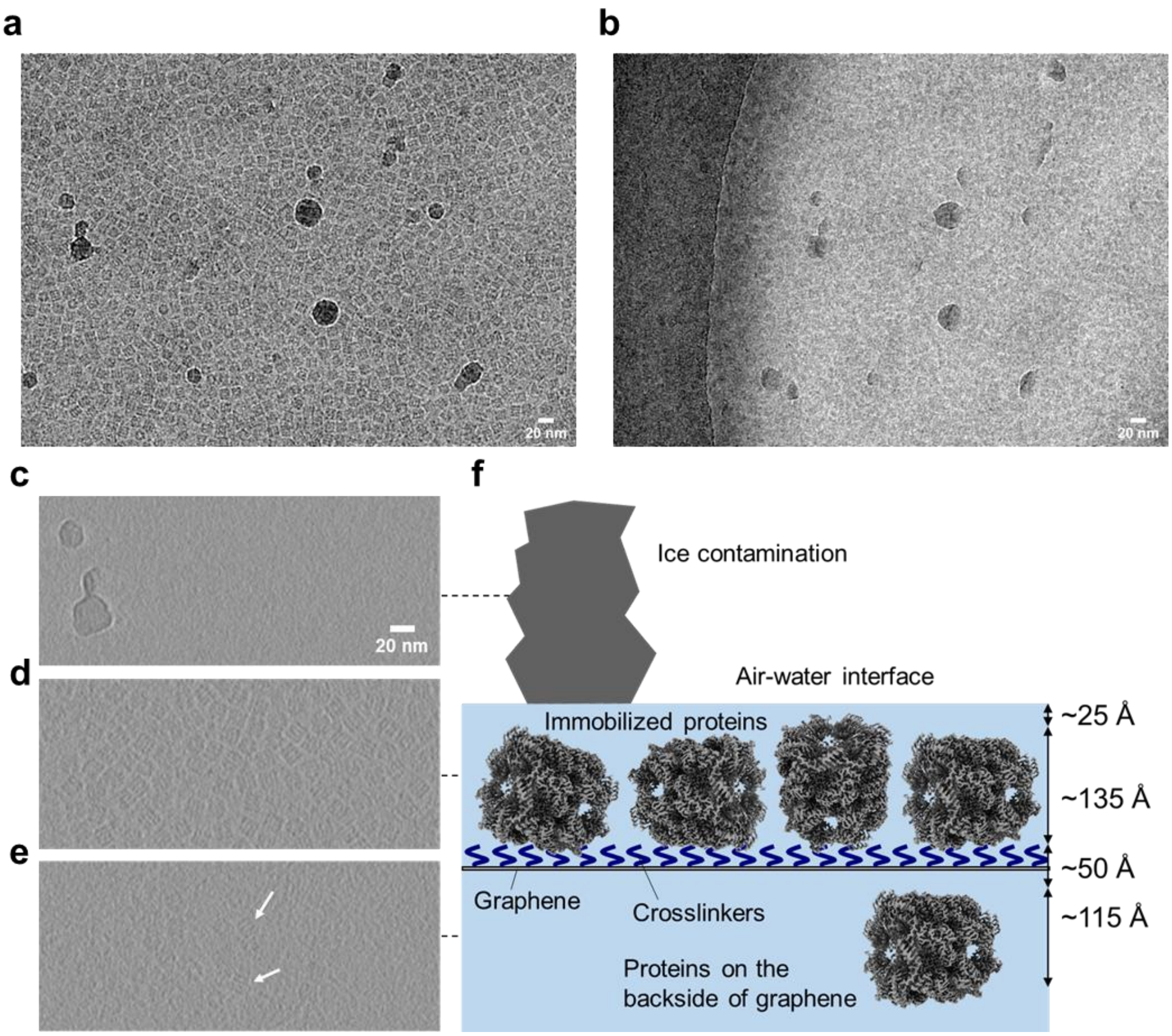


**Supplementary Figure 5. ECT of GroEL-embedded ice film on the EG-grid. a**,**b**, CryoEM images of GroEL-embedded ice on the EG-grid at a tilt angles of 0º (**a**) and –54º (**b**). **c-e**, Three layer images extracted from the tomogram reconstructed from the area corresponding to those shown in (**a**) and (**b**). (**c**) A layer near the surface of vitreous ice where the same pair of ice contaminants seen in the left part of (**a**) and (**b**) can be seen. (**d**) A vitreous ice layer containing GroEL particles, which is ~240 Å lower than that of (**c**). (**e**) Another layer showing a few GroEL particles (white arrows), which is ~225 Å lower than that of (**d**). See the whole tomogram in Supplementary Video 2. **f**, Schematic illustration of ice-embedded GroEL particles attached on the graphene surface of the EG-grid.

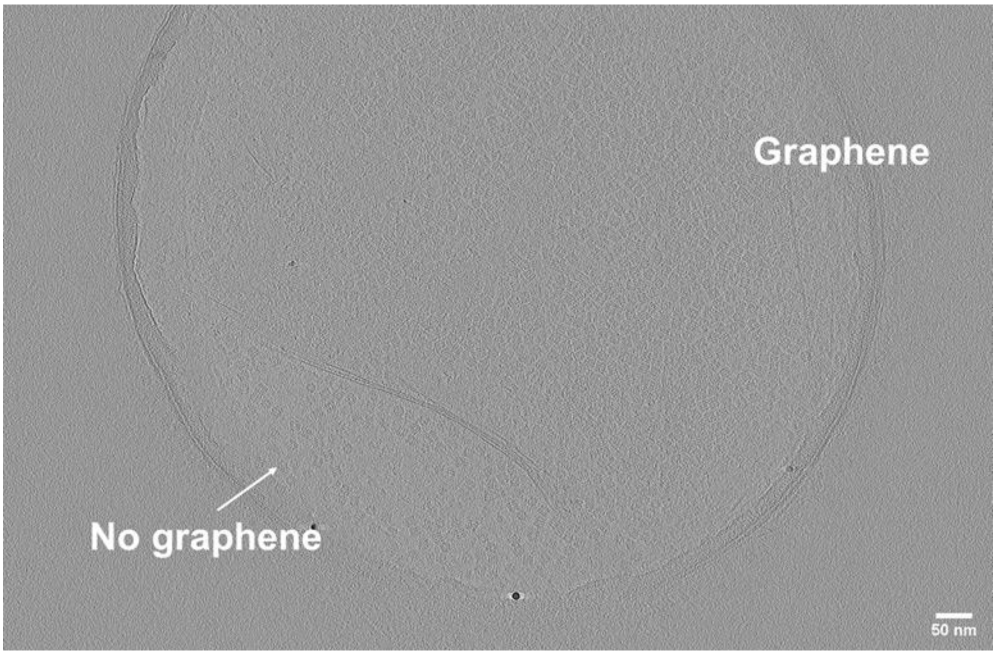


**Supplementary Figure 6. ECT of GroEL-embedded ice film on the glow discharged graphene grid.** A layer image extracted from the reconstructed tomogram. The hole included both area with and without graphene. See the whole tomogram in Supplementary Video 3.

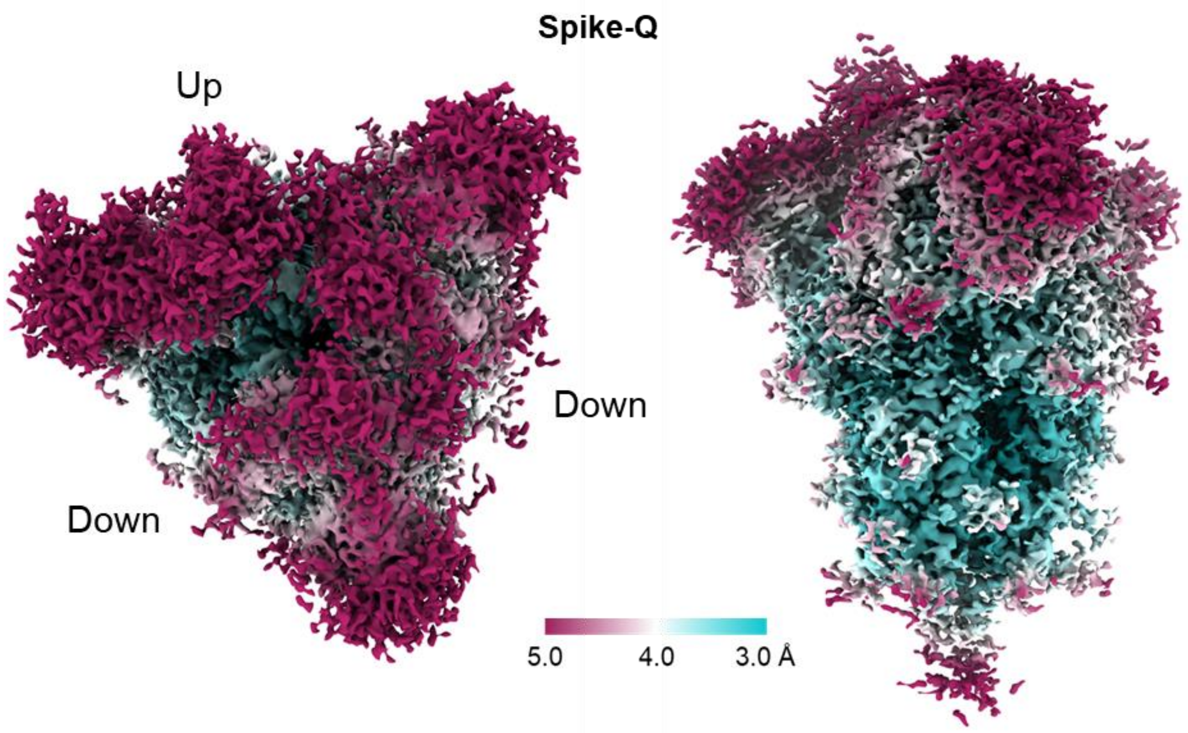


**Supplementary Figure 7. CryoEM image analysis of the Spike-Q dataset.** Final 3D map of the spike-Q dataset in two orthogonal views: top view, left panel; and side view, right panel. The local resolution distribution is colored as in the color bar.

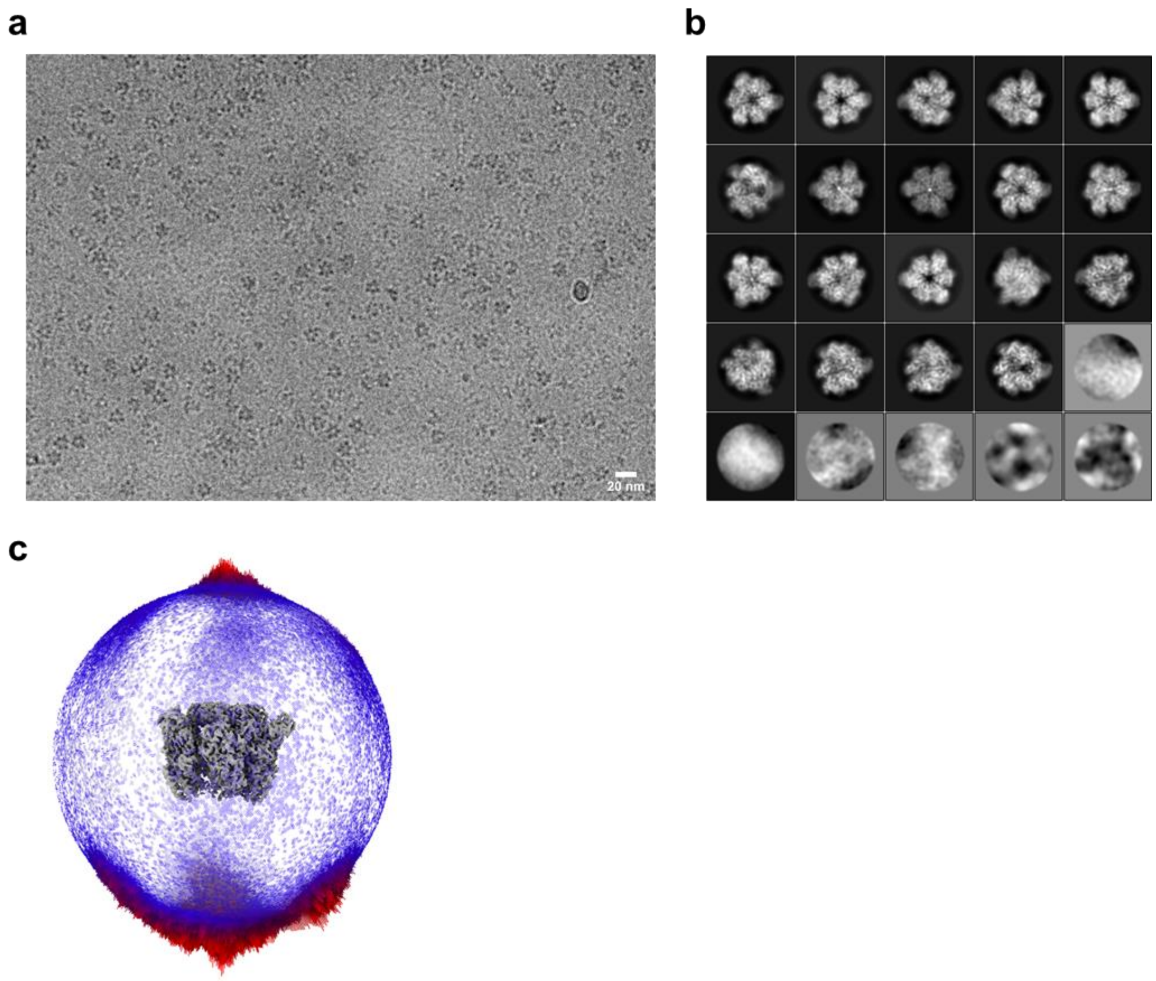


**Supplementary Figure 8. CryoEM image analysis of the $V_1$-ATPase-EG dataset. a**, Typical cryoEM image (50,000 x) of the $A_3B_3$ ring of $V_1$-ATPase on the EG-grid. **b**, Top 25 2D class averages aligned in the descending order of particle numbers from left to right and top to bottom. **c**, Angular distribution of particles used in the final refinement. The final 3D map is also shown in grey for reference.

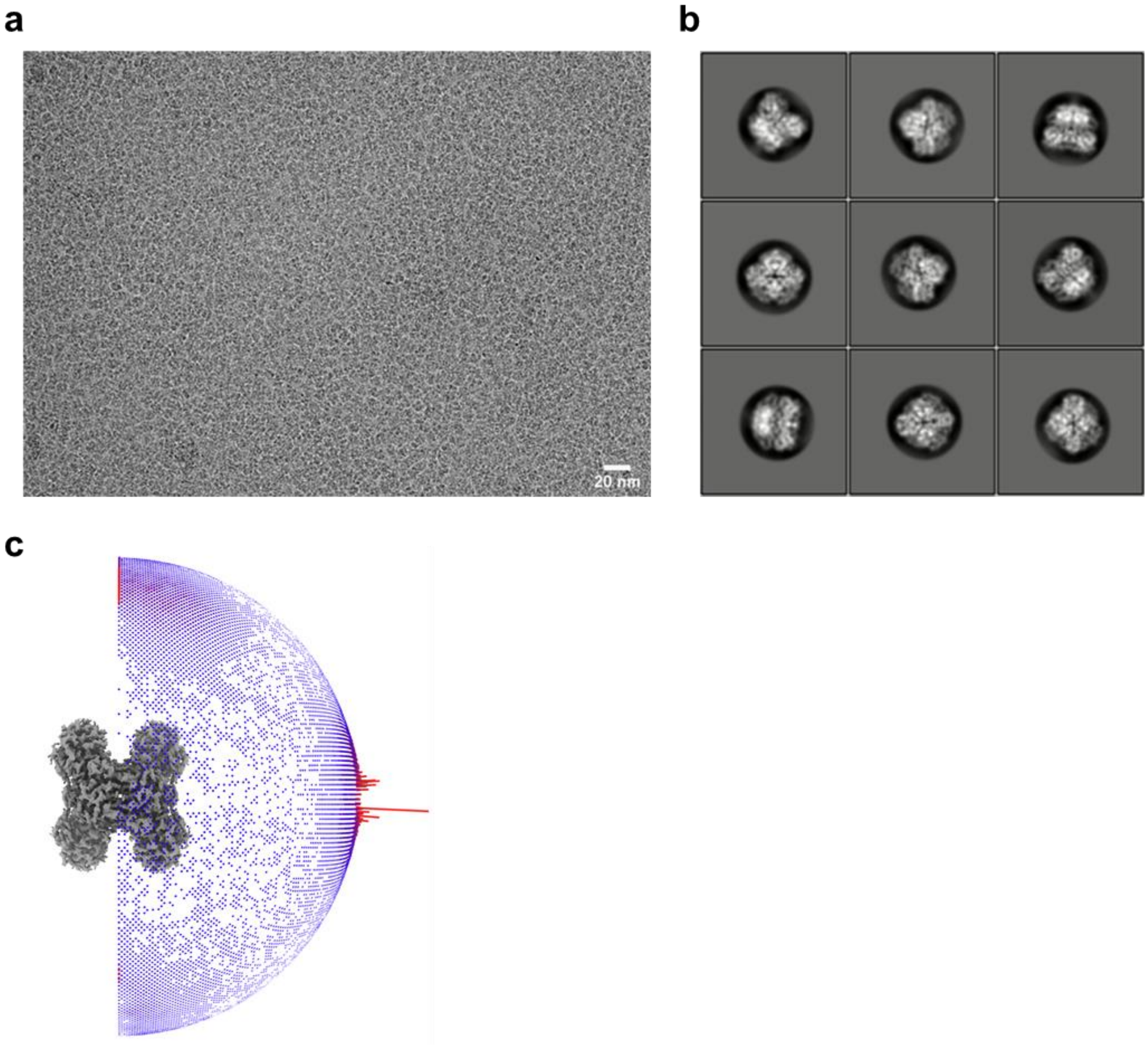


**Supplementary Figure 9. CryoEM image analysis of the GAPDH-EG dataset. a**, Typical cryoEM image (60,000 x) of GAPDH on the EG-grid. **b**, Top 9 2D class averages aligned in the descending order of particle numbers from left to right and top to bottom. **c**, Angular distribution of particles used in the final refinement. The final 3D map is also shown in grey for reference.

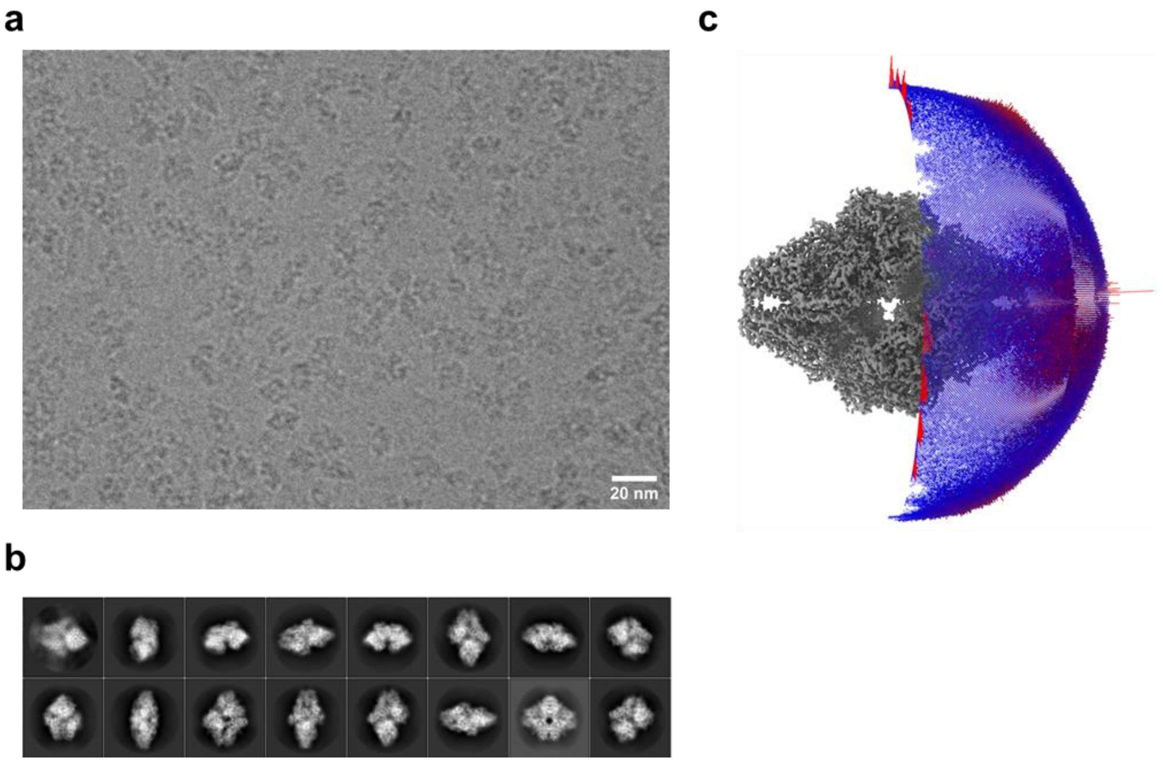


**Supplementary Figure 10. CryoEM image analysis of the β-galactosidase-EG dataset. a**, Typical cryoEM image (100,000 x) of β-galactosidase on the EG-grid. **b**, Top 16 2D class averages aligned in the descending order of particle numbers from left to right and top to bottom. **c**, Angular distribution of particles used in the final refinement. The final 3D map is also shown in grey for reference.

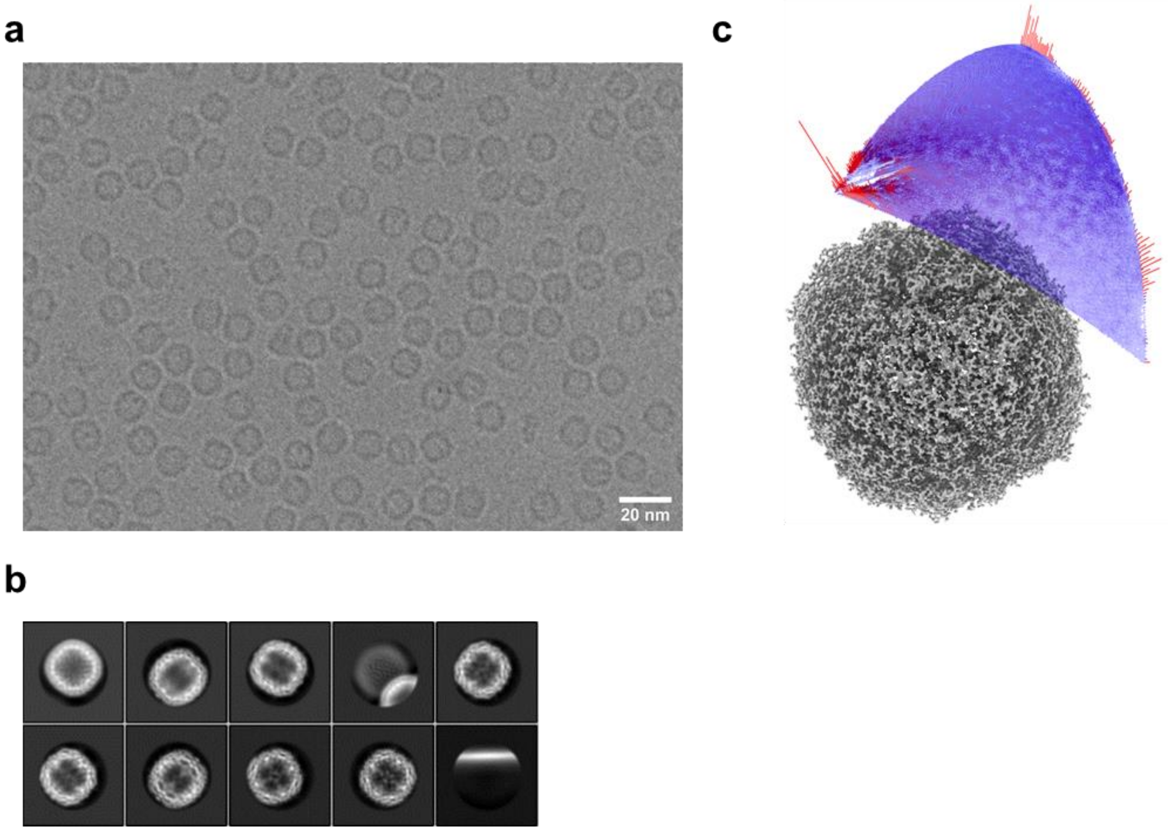


**Supplementary Figure 11. CryoEM image analysis of the apoferritin-EG dataset. a**, Typical cryoEM image (120,000 x) of apoferritin on the EG-grid. **b**, Top 10 2D class averages aligned in the descending order of particle numbers from left to right and top to bottom. **c**, Angular distribution of particles used in the final refinement. The final 3D map is also shown in grey for reference.

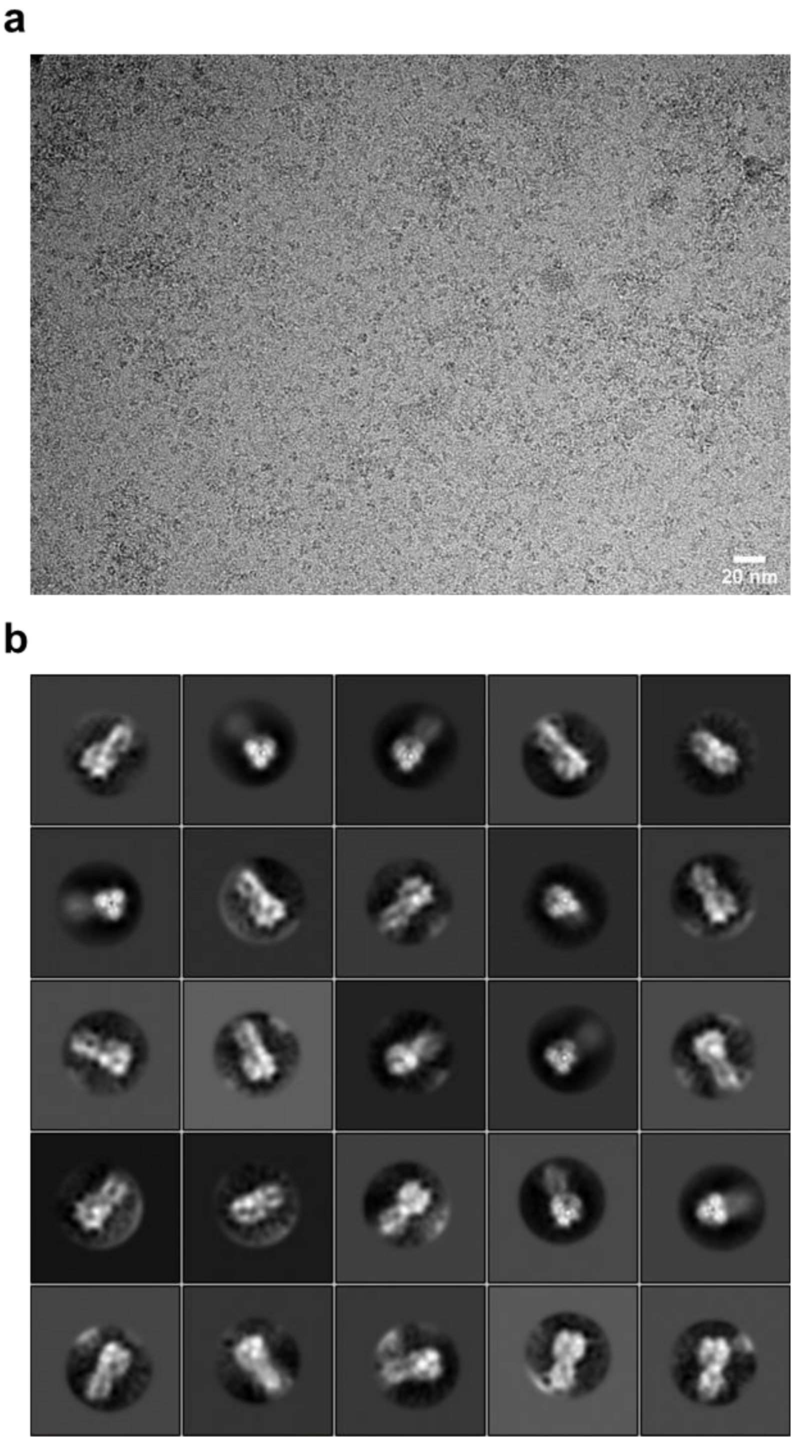


**Supplementary Figure 12. CryoEM image analysis of the hemagglutinin on EG-grid. a**, Typical cryoEM image (60,000 x) of hemagglutinin on the EG-grid. **b**, Top 25 2D class averages aligned in the descending order of particle numbers from left to right and top to bottom.

### Supplementary Table 1. CryoEM data collection and image processing.

| Dataset | GroEL-EG (EMDB-31310) | GroEL-Gl (EMDB-31311) | GroEL-Gh (EMDB-32159) | SARS-CoV-2 Spike-EG (EMDB-32160) | SARS-CoV-2 Spike-Q (EMDB-32161) |
|---|---|---|---|---|---|
| Magnification | 60,000 | 60,000 | 60,000 | 60,000 | 60,000 |
| Voltage (kV) | 300 | 300 | 300 | 300 | 300 |
| Electron exposure ($e^{-}/Å^{2}$) | 40 | 40 | 40 | 60 | 60 |
| Defocus range (μm) | −0.5 to −2.0 | −0.5 to −2.0 | −0.5 to −2.0 | −0.5 to −2.0 | −0.5 to −2.0 |
| Pixel size (Å) | 0.813 | 0.816 | 0.878 | 0.870 | 0.870 |
| Symmetry imposed | D7 | D7 | D7 | C1 | C1 |
| Initial images (no.) | 504 | 5,614 | 649 | 1,163 | 1,029 |
| Optics group (no.) | 10 | 11 | 13 | 12 | 10 |
| Initial particle images (no.) | 178,215 | 1,118,169 | 340,446 | 602,489 | 244,439 |
| Final particle images (no.) | 158,485 | 215,011 | 198,677 | 150,316 | 58,102 |
| Final/Initial particle ratio (%) | 88.9 | 19.2 | 58.4 | 24.9 | 23.8 |
| Average of final particles per image (no.) | 314.5 | 38.3 | 306.1 | 129.2 | 56.5 |
| Map resolution (Å) | 1.99 | 2.08 | 2.09 | 3.10 | 3.23 |
| FSC threshold | 0.143 | 0.143 | 0.143 | 0.143 | 0.143 |
| Map resolution range (Å) | 1.93–3.04 | 2.02–3.09 | 2.06–3.07 | 2.82–14.7 | 2.95–12.2 |

| Dataset | $V_1$-ATPase-EG (EMDB-31312) | GAPDH-EG (EMDB-32162) | β-gal-EG (EMDB-31313) | Apoferritin-EG (EMDB-31314) |
|---|---|---|---|---|
| Magnification | 50,000 | 60,000 | 100,000 | 120,000 |
| Voltage (kV) | 300 | 300 | 300 | 300 |
| Electron exposure ($e^{-}/Å^{2}$) | 62 | 40 | 40 | 40 |
| Defocus range (μm) | −1.0 to −2.5 | −0.5 to −2.0 | −0.5 to −2.0 | −0.3 to −1.3 |
| Pixel size (Å) | 0.990 | 0.813 | 0.495 | 0.490 |
| Symmetry imposed | C1 | D2 | D2 | O |
| Initial images (no.) | 2,052 | 241 | 3,242 | 7,500 |
| Optics group (no.) | 11 | 8 | 16 | 12 |
| Initial particle images (no.) | 1,193,032 | 301,451 | 491,997 | 793,398 |
| Final particle images (no.) | 238,765 | 88,731 | 231,395 | 527,261 |
| Final/Initial particle ratio (%) | 20.0 | 29.4 | 47.0 | 66.5 |
| Average of final particles per image (no.) | 116.4 | 368.2 | 71.4 | 70.3 |
| Map resolution (Å) | 3.03 | 2.16 | 1.81 | 1.29 |
| FSC threshold | 0.143 | 0.143 | 0.143 | 0.143 |
| Map resolution range (Å) | 2.85–4.66 | 2.10–2.98 | 1.76–2.35 | 1.28–1.47 |

**Supplementary Video 1. Preparation of the EG-grid.**

**Supplementary Video 2. Reconstructed tomogram of GroEL on the EG-grid.** The section starts from near the surface of vitreous ice, goes through the GroEL and graphene layer on the EG-grid and then goes backward. Tomogram thickness is 100 nm.

**Supplementary Video 3. Reconstructed tomogram of GroEL on the glow discharged graphene grid.** The section starts from near the surface of vitreous ice, goes through the GroEL and graphene layer and then goes backward. Tomogram thickness is 100 nm.
